# The Cost of Untracked Diversity in Brain-Imaging Prediction

**DOI:** 10.1101/2021.06.16.448764

**Authors:** Oualid Benkarim, Casey Paquola, Bo-yong Park, Valeria Kebets, Seok-Jun Hong, Reinder Vos de Wael, Shaoshi Zhang, B.T. Thomas Yeo, Michael Eickenberg, Tian Ge, Jean-Baptiste Poline, Boris Bernhardt, Danilo Bzdok

## Abstract

Brain-imaging research enjoys increasing adoption of supervised machine learning for single-subject disease classification. Yet, the success of these algorithms likely depends on population diversity, including demographic differences and other factors that may be outside of primary scientific interest. Here, we capitalize on *propensity scores* as a composite confound index to quantify diversity due to major sources of population stratification. We delineate the impact of population heterogeneity on the predictive accuracy and pattern stability in two separate clinical cohorts: the Autism Brain Imaging Data Exchange (ABIDE, n=297) and the Healthy Brain Network (HBN, n=551). Across various analysis scenarios, our results uncover the extent to which cross-validated prediction performances are interlocked with diversity. The instability of extracted brain patterns attributable to diversity is located preferentially to the default mode network. Our collective findings highlight the limitations of prevailing deconfounding practices in mitigating the full consequences of population diversity.

## Introduction

Brain scanning technology opens a non-invasive window into the structure and function of the human brain. Combined with machine learning algorithms, brain-imaging research is now gaining momentum to transition from group-level contrast analyses towards single-subject prediction (Bzdok, 2017; Bzdok & Yeo, 2017; Gabrieli et al., 2015). In supervised learning, the main purpose is to learn coherent patterns from brain measurements (i.e., brain signatures) that can be used to make accurate forecasts for new subjects (Orru et al., 2012; Pereira et al., 2009). The prediction paradigm holds the promise of improving disease diagnosis, enhancing prognostic estimates, and ultimately paving the way to precision medicine (Brammer, 2009; Bzdok et al., 2021). Machine learning methods are now increasingly adopted for the goal of classifying various conditions, including autism spectrum disorder (ASD) (Heinsfeld et al., 2018; Plitt et al., 2015; Sabuncu et al., 2015; Wolfers et al., 2019), attention-deficit/hyperactivity disorder (ADHD) (Brown et al., 2012; Riaz et al., 2020; Sen et al., 2018; Wang et al., 2018), anxiety (ANX) (Frick et al., 2014; Liu et al., 2015), or schizophrenia (Davatzikos et al., 2005; Karrer et al., 2019; Rozycki et al., 2018; Shen et al., 2010; Yassin et al., 2020). However, there is a large variability in the accuracy reports across studies (Arbabshirani et al., 2017; Pulini et al., 2019; Woo et al., 2017). Counterintuitively, the larger the clinical cohorts have become, the worse the prediction performance of machine learning algorithms has become. The unknown reasons behind this varying success hamper the translation of emerging neuroimaging findings into clinical practice.

In the current landscape of machine learning applications in neuroimaging, researchers are facing challenges related to the generalizability and replicability of brain patterns valuable for prediction. Careful validation of machine learning tools is therefore needed to ascertain the robustness of the learned brain patterns. When using small and homogeneous datasets, however, common crossvalidation procedures are more likely to yield high prediction accuracies, although at the expense of producing biased biomarkers with poor generalization to new or future cohorts. The last two decades have witnessed an unprecedented growth in the number of open-access data-sharing initiatives in the neuroscientific community (Alexander et al., 2017; Biswal et al., 2010; Casey et al., 2018; Di Martino et al., 2014, 2017; Van Essen et al., 2013). In these initiatives, data are collected from multiple acquisition sites to aggregate larger and demographically more heterogeneous datasets. Such multi-site data pooling efforts are more likely to reflect the true diversity in the wider populations. This first wave of retrospective data collection initiatives offers great opportunities for the advancement of scientific discoveries in neuroscience (Milham et al., 2018; Poldrack et al., 2017).

However, these efforts also pose new hurdles due to the diverging characteristics and backgrounds of the subjects who are recruited at each site. Discrepancies in demographic and clinical variables may interfere with the identification of robust relationships between imaging-based features and the target variables (e.g., diagnosis). These challenges may entail potential bias for downstream decision-making by healthcare professionals and other stakeholders. Moreover, gathering brain scans from multiple sites comes with the potential cost of increased heterogeneity due to different sources of variation, such as batch effects (Kostro et al., 2014; Yamashita et al., 2019). Compared to cohorts recruited from a single site, classification in multi-site cohorts has shown lower performance and often poor generalization to subjects from new scanning sites (Arbabshirani et al., 2017; Chen et al., 2016; Nielsen et al., 2013; Schnack & Kahn, 2016). When handled with insufficient care, these general sources of variation that tell people apart may prevent predictive models from learning accurate patterns and weaken their performance. For these reasons, faithful disease classification in multi-site cohorts may increasingly require sharp tools that take several dimensions of diversity into account.

Previous studies have typically focused on assessing generalizability along a single dimension: for example, evaluating a trained predictive model on a specific subset of subjects from unseen scanning sites (e.g., leave-N-sites-out), a subset of subjects with a specific sex or age range (e.g., Abraham et al., 2017; Bonkhoff et al., 2021; Kiesow et al., 2021; Lanka et al., 2019). The increasingly embraced crossvalidation tactics can monitor and probe the effects from one specific source of heterogeneity. Yet, cohort heterogeneity may stem from multiple aspects of diversity (e.g., age, sex and acquisition site). For example, a performance decay in held-out acquisition sites might be in part attributable to other aspects of population stratification (e.g., age and sex distribution). To deal with such diversity, we need approaches that can appropriately incorporate sources of population heterogeneity that should be acknowledged in the analytical workflow (cf. Bzdok et al., 2020).

Our study aims to provide investigators with a handle to directly quantify the ramifications of population diversity. We propose to recast the propensity score framework to faithfully track out-of-distribution behavior of predictive analysis pipelines. Briefly, the propensity score has originally been introduced for computing the probability of treatment assignment given an individual’s collective covariates (Rosenbaum & Rubin, 1983). An appealing property of propensity scores is the opportunity to encapsulate a mixed envelope of covariates and express them as a single dimension of stratification. Once built, a propensity score model can be deployed for a variety of purposes, including matching, stratification, covariate adjustment and weighting (Ali et al., 2019; Austin, 2011a). Henceforth, our notion of diversity relies on continuous differences in the propensity scores of the subjects in a cohort. Subjects who are closer to each other on the propensity score spectrum can be assumed to have more similar constellations of covariates. Larger differences in propensity scores indicate bigger gaps in population stratification or background between a given pair of subjects. After stratifying the cohort by means of propensity scores, we examined different sampling schemes to analyze the impact of diversity on the performance of our predictive models and assess their generalizability. For the estimation of propensity scores and thus diversity, we considered a set of non-imaging covariates with potential confounding effects that are widely used and readily available in most neuroimaging datasets (Dinga et al., 2020; Rao et al., 2017; Wachinger et al., 2020): age, sex assigned at birth, and scanning site. Results on the classification of subjects with a neurodevelopmental diagnosis versus healthy controls from both the Autism Brain Imaging Data Exchange (ABIDE) and Healthy Brain Network (HBN) cohorts using functional and structural brain measurements show that, even after rigorous matching and nuisance deconfounding, diversity exerts a substantial impact on the generalization accuracy of the predictive models and on the pattern stability of the extracted biomarkers.

## Methods

### Participants and brain scanning resources

We examined resting-state functional MRI (rs-fMRI) and cortical thickness data from both waves of the openly-shared Autism Brain Imaging Data Exchange initiative (ABIDE I and II; http://fcon_1000.projects.nitrc.org/indi/abide) (Di Martino et al., 2014, 2017) and the Healthy Brain Network (HBN) dataset (Alexander et al., 2017), releases 1-8 collected between June 2017 and January 2020.

For ABIDE, we considered data from all acquisition sites with at least 10 subjects per group and with both children and adults. After detailed quality control, only cases with acceptable T1-weighted (T1w) MRI, surface-extraction, and head motion in rs-fMRI were included. Subjects with ASD were diagnosed based on an in-person interview administered by board-certified mental health professionals using the gold-standard diagnostics of the Autism Diagnostic Observation Schedule, ADOS (Lord et al., 1999) and/or Autism Diagnostic Interview-Revised (ADI-R) (Lefort-Besnard et al., 2020; Lord et al., 1994). Typically-developing (TD) subjects had no history of mental disorders. For all groups, subjects were excluded on the basis of genetic disorders associated with autism (i.e., Fragile X), contraindications to MRI scanning, or current pregnancy. The ABIDE data collections were performed in accordance with local Institutional Review Board guidelines, and data were fully anonymized. These quality procedures resulted in a total of 297 subjects (151 ASD and 146 TD) from 4 different sites (i.e., NYU, PITT, USM, TCD) in ABIDE.

For HBN, we included 102 TD individuals and 449 subjects carrying a diagnosis of ASD, ADHD and/or anxiety (ANX). From all subjects with at least one diagnosis, 53 were diagnosed with ASD and ADHD, 12 diagnosed with ASD and ANX, 99 with ANX and ADHD, and 17 subjects were diagnosed with all three. Only 24 ASD, 171 ADHD and 73 ANX subjects had no comorbidity. The HBN cohort offers a broad range of heterogeneity in developmental psychopathology. Participants were recruited using a community-referral model (for inclusion criteria, see https://fcon_1000.projects.nitrc.org/indi/cmi_healthy_brain_network/Recruitement.html#inclusion-exclusion). Psychiatric diagnoses were assessed and reported by clinicians according to DSM-5 criteria (APA, 2013). HBN was approved by the Chesapeake Institutional Review Board. Written informed consent was obtained from all subjects and from legal guardians of participants younger than 18 years. Hence, after assessing the same criteria for quality control, we ended up with a total of 551 subjects, with several subjects having more than one neurodevelopmental diagnosis, from 3 different sites (i.e., SI, RU, CBIC) in HBN.

More details about acquisition settings and demographic information about the subjects included in our study from both datasets are reported in **Tables S1 and S2**, respectively.

### Brain scan preprocessing workflow

For both ABIDE and HBN datasets, structural brain scans (T1w MRI) were preprocessed with FreeSurfer (Dale et al., 1999; Fischl, 2012; Fischl et al., 1999). The pipeline performed automated bias field correction, registration to stereotaxic space, intensity normalization, skull-stripping, and tissue segmentation. White and pial surfaces were reconstructed using triangular surface tessellation and topology-corrected. Surfaces were inflated and spherically registered to the fsaverage5 template. Segmentations and surfaces were subject to visual inspection. Subjects with erroneous segmentations were excluded from further analysis.

For the resting-state functional brain scans in ABIDE, we built on data previously made available by the Preprocessed Connectomes initiative (http://preprocessed-connectomes-project.org/abide). The preprocessing was performed with C-PAC (https://fcp-indi.github.io) and included slice-time correction, head motion correction, skull stripping, and intensity normalization. The rs-fMRI data were de-trended and adjusted for common nuisance effects related to head motion, white matter and cerebrospinal fluid signals using CompCor (Behzadi et al., 2007), followed by band-pass filtering (0.01-0.1 Hz). For the rs-fMRI data in HBN, we discarded the first five volumes, removed the skull, and corrected for head motion. Magnetic field inhomogeneity was corrected using topup with reversed phase-encoded data (Andersson et al., 2003). After applying a high-pass filter at 0.01 Hz, nuisance effects were removed using ICA-FIX (Salimi-Khorshidi et al., 2014). We excluded subjects who had mean framewise displacement greater than 0.3. Individual rs-fMRI data were mapped to the corresponding mid-thickness surfaces, resampled to the Conte69 template (https://github.com/Washington-University/Pipelines), and smoothed using a 5 mm full-width-at-half-maximum (FWHM) kernel.

### Pattern-learning pipeline for disease classification

The construction of the feature space from the brain scans was based on a topographical brain parcellation: 100 functionally-defined cortical regions from a widely used reference atlas (Schaefer et al., 2018). For cortical thickness, each subject’s fingerprint consisted of a 100-feature vector corresponding to the mean cortical thickness of the vertices within each of the 100 atlas regions. Similarly, for intrinsic functional fluctuations, the time series were averaged over all vertices in each of the 100 atlas regions and used to build functional connectivity profiles based on all pairwise correlations (Pearson’s correlation coefficient) between the 100 region time series. For the rs-fMRI, the feature vectors thus consisted of 4950 unique region-region connectivity strengths.

The predictive model we used for classification (e.g., ASD vs TD) is the commonly used logistic regression model with an optimization loss that includes a penalty term for Tikhonov (*l*_2_) regularization. Let *w* be the coefficients (i.e., weights) of our model and *x* the feature vector (e.g., of cortical thickness values) corresponding to a given subject. For prediction of continuous class assignments in form of log odds, the model specification involves mapping the weighted combination of features (i.e., *z* = *w^T^x*) from input space (cf. feature engineering above) into a probability using the sigmoid function. For gradient-descent-based estimation of this convex optimization problem, the set of final model coefficients indicating the global minimum was obtained by solving a structured risk minimization problem based on the following cost function:

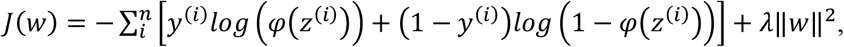

where *n* is the number of observations, *y*^(*i*)^ carries the class label (i.e., ASD or TD), *z*^(*i*)^ is the weighted combination of features of the *i*-th observation, and *φ* denotes the sigmoid function, which is defined as *φ*(*z*) = 1/(1 + *e^−z^*). The first term is the log-likelihood function, and the second term penalizes the weight coefficients, with *λ* denoting the hyperparameter that controls the regularization strength of the *l*_2_ penalty term. Tikhonov regularization is known to exert smooth parameter shrinkage by acting on the low-variation directions that underlie the input space relatively more than the dominant high-variation patterns in the data (Bishop, 2006). This model has been shown to perform better than several competing machine learning models in comprehensive benchmark studies of ASD classification (Abraham et al., 2017) and other commonly studied phenotype predictions (Kiesow et al., 2021; Schulz et al., 2020).

To rigorously evaluate our analytical workflow, we adopted a nested cross-validation (CV) strategy. Regardless of how the outer CV was implemented (see paragraph below on *out-of-distribution prediction*), we consistently used an inner CV based on 5 folds for tuning the hyperparameter controlling the regularization strength (i.e., *λ*). The value for the complexity parameter *λ* was chosen from a grid of 7 equidistant points in logarithmic scale in the interval [1e-3, 1e+3]. Prior to fitting our model to the data, brain features were z-scored across subjects on the training set at hand (i.e., respecting the mechanics of cross-validation), and the derived parameters were then used to apply z-scoring of the features to the subjects in the test set.

### Quantifying diversity using propensity scores

Our overarching aim was to characterize the role of population heterogeneity in forming prediction by means of pattern extraction. The notion of diversity used throughout this work leans on three important facets of population stratification that are available in most population datasets: subject age, subject sex and scanning site. In our study, we proposed to get a handle on this notion of subject diversity by virtue of the propensity score framework that is established in a variety of domains (Rosenbaum & Rubin, 1983). Diversity thus denoted the distance based on these common covariates between subjects in our cohorts. Observational studies have benefited from propensity scores to successfully estimate the probability of a subject receiving treatment given the covariates. Here, we concentrated on estimating the probability that a given subject carries a disease diagnosis based on a set of covariates, that is:

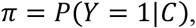

where *Y* is the target variable, with *Y* = 1 for ASD and *Y* = 0 for TD. *C* are the covariates, which collectively provided the basis for quantifying population diversity. In other words, we estimated a model for the purpose of deriving a propensity score as a function of the covariates of a given subject. Although other machine learning tools have sometimes been used to obtain propensity scores (Lee et al., 2011; Setoguchi et al., 2008; Westreich et al., 2010), logistic regression is probably the most common implementation in this context. This natural choice of method yields the following model specification for our propensity scores framework:

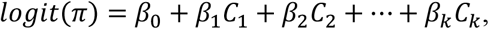

where *k* denotes the number of covariates or sources of population stratification, *β*_0_ is the intercept capturing the average classification log odds in the subject data at hand, while *logit*(·) is a non-linear transformation (called ‘link function’ in the generalized linear modeling family) that maps from probability space to log odds space. In other words, we invoked a conditional expectation that is implemented by a logistic regression model to predict diagnosis (e.g., ASD) using the covariates of each subject as features. Although propensity scores are often based on treatment assignment, for simplicity, we use them throughout this work to refer to the probability we defined above. We emphasize that the propensity model itself does not involve or have access to any of the brain-imaging features of primary scientific interest.

A key asset of the propensity score framework is the opportunity to seamlessly construct subsets of subjects with homogeneous background profiles with respect to the observed covariates. This follows because the propensity score is a balancing score. At each value *p* of the propensity score, the distributions of the covariates are the same in both groups (Rosenbaum & Rubin, 1983; Stuart, 2010), that is:

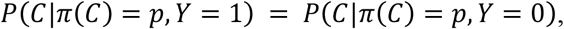

where *π*(*C*) denotes the propensity score corresponding to the covariate set *C*, and Y is a binary variable holding the diagnosis.

Multiple applications of propensity scores can be used for confounding adjustment including matching, stratification and inverse probability weighting (Ali et al., 2019; Austin, 2011a). In previous neuroimaging studies, propensity scores have been used to address confounding in permutation testing (Hedlin et al., 2010), classification (Linn et al., 2016) and regression (Rao et al., 2017) problems. An appealing quality of propensity scores is a form of dimensionality reduction: representing a mixed envelope of covariates as a single number that coherently encapsulates the sources of inter-subject variation indexed by the joint covariates. Matching, for example, can then be performed directly on the propensity scores rather than on the covariates, which rapidly becomes infeasible with high-dimensional vectors of covariates. Moreover, propensity scores can be readily used to handle both continuous (e.g., age) and categorical (e.g., site or sex) covariates in a single coherent framework. Any possible collinearity in coefficient estimation of the propensity score model does not affect the results because this modeling step follows the prediction goal (Bzdok & Ioannidis, 2019). In other words, we are not concerned with point estimate, interval estimate or subject-matter interpretation of any specific coefficient inside the propensity score model itself (McMurry et al., 2015).

After estimating the propensity scores, subjects were spanned out along a one-dimensional continuum that gauges the diversity among them: the more dissimilar the subjects in their covariates, the more distant they were from each other on the propensity score spectrum. By chunking the subjects into strata with similar confounding architecture, within each of which subjects had minimal diversity relative to each other, we could assess the generalizability of our predictive models sampling different strata for train and test in an operationalizable series of experiments.

### Matching and stratification

Naively partitioning the data points however may produce strata with unbalanced classes. To safeguard against skewing of results due to class imbalance, we initially matched subjects based on their propensity score estimates. We employed a pair matching approach: one specific subject in the TD group was allocated to one subject in the ASD group. To identify these pairs in a data-driven fashion, we carried out the matching procedure using the Hungarian algorithm (Kuhn, 1955). The Hungarian algorithm obtained a globally optimal pairing by minimizing the global distance (i.e., a ‘cost’ quantity) between paired subjects based on the absolute difference of their estimated propensity scores. Nonetheless, this approach is not exhaustive, in that, the absolute differences between matched subjects are unlikely to be zero. This circumstance especially comes into play when there is imperfect overlap between the overall distributions of estimated propensity scores in ASD and TD. This is because there is a risk of ending up with paired subjects with very dissimilar propensity scores, and hence different positioning according to directions of population stratification.

To overcome this hurdle, we added a restriction to the Hungarian algorithm such that the absolute difference in the propensity scores of matched subjects must not exceed a pre-specified threshold - the so-called caliper (Lunt, 2014). This constraint is commonly achieved by setting the cost of matching pairs of subjects to a large value when their difference exceeds the threshold. Then, paired subjects returned by the Hungarian algorithm are inspected to discard those pairs that exceed the threshold. The value must be larger than all values in the cost matrix of the Hungarian algorithm. In this work, the caliper was set to 0.2, which is a popular choice in the literature (Austin, 2011b; Wang et al., 2013). The maximum acceptable distance for pairs of subjects to be considered a match was then set to 0.2 * *d_sd_*, where *d_sd_* was the standard deviation of all pairwise absolute differences between the subjects in our dataset. The benefit of using a caliper was that we achieved a better matching of the considered subjects, at the possible expense of reducing the number of subjects in a data analysis scenario. Thus, final pairs of TD-ASD subjects had very similar covariates. As shown in **Figure 1A**, distributions of propensity scores after matching showed better overlap (middle panel) than the unmatched original distributions (left panel), especially in HBN.

**Figure 1.**
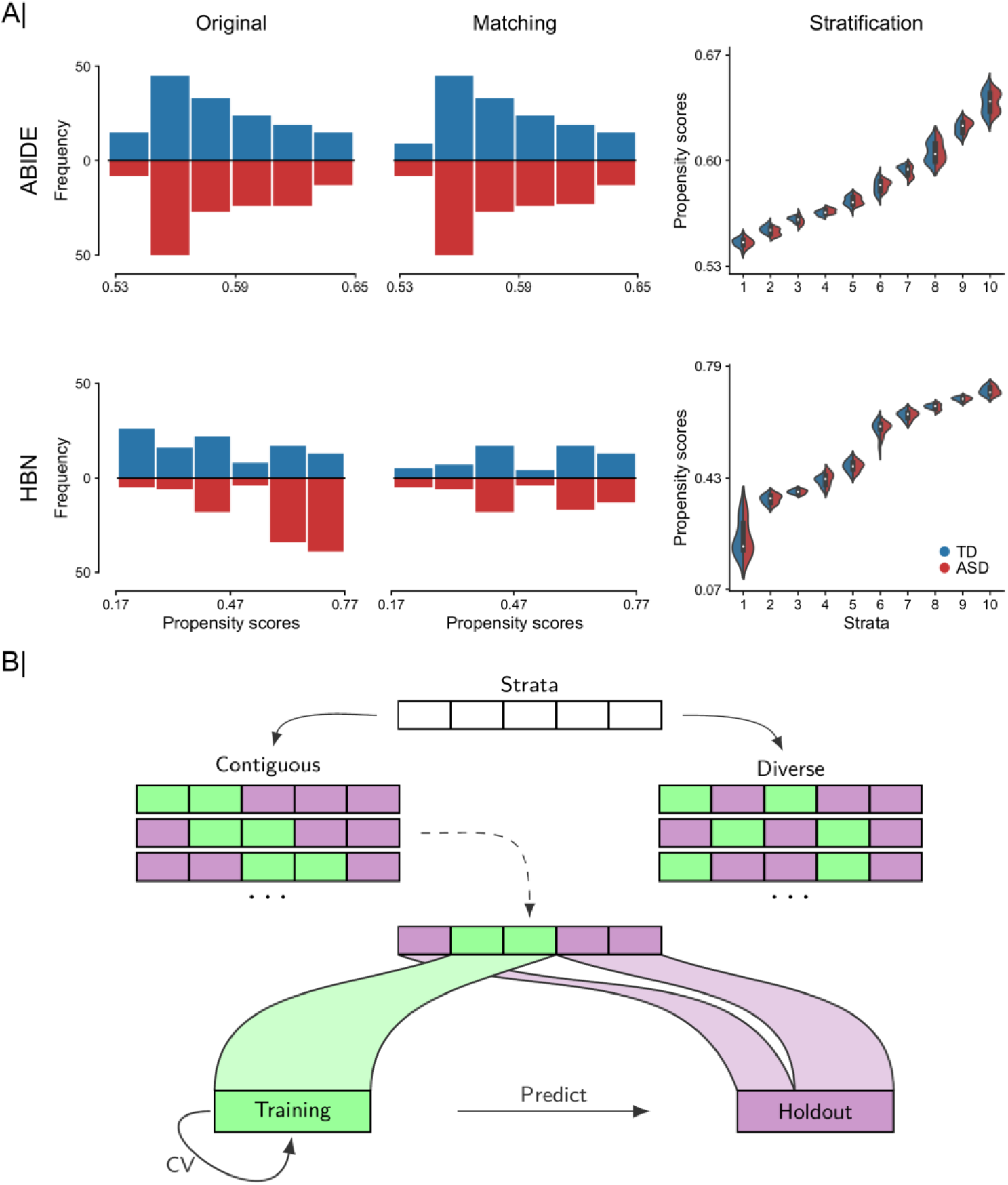
Workflow for systematic subject stratification and diversity-based subject sampling. We repurpose propensity scores as a tool to translate several different indicators of population stratification into a single diversity index. A) From left to right: two-sided histograms in which we show bin counts bins of estimated propensity score distributions for typically developing (TD, blue) and autism spectrum disorder (ASD, red) subjects before (i.e., using all observations in original dataset) and after matching (based on age, sex, site), and partitioning of matched subjects in 10 equally-sized subject sets (i.e., strata) based on similarities of their propensity scores for ABIDE (top) and Healthy Brain Network (HBN; bottom) cohorts. Distributions of propensity scores after matching showed better overlap (middle panel) than the unmatched original distributions (left panel), especially in HBN. Propensity scores are computed as a function of age, sex and site, which are commonly available in many future population cohorts. B) Diversity-based subject sampling: Given the attribution of each subject to 1 of *q* homogeneous strata, a subset of *r* strata is picked and combined to form the training set used for estimating the predictive model (green) and the *q*-*r* remaining strata served as held-out set for testing the performance of that learning model (violet). Two different sampling regimes are used to form the training set: *i)* a *contiguous* scheme where the training set is composed of adjacent strata of similar diversity, and *ii)* a *diverse* scheme where the training data is composed of non-contiguous strata (at least one) with subjects pulled from diverse populations. For each analysis setting, the classification accuracy is assessed within the training set based on rigorous 10-fold fit-tune-predict cross-validation (CV) cycles, and then all the training data is used to predict disease status in unseen holdout subjects.

After the matching step, the aligned subject pairs were ranked according to their mean estimated propensity scores (i.e., the mean propensity score of a matched pair of subjects) and stratified into mutually exclusive subsets, henceforth ‘strata’. In our work, subjects were divided into 10 equally-sized strata using the deciles of the estimated propensity scores. Although matching based on the mean propensity score of paired subjects may not be perfect in all cases, **Figure 1A** shows good overlap between the distributions of propensity scores for each group (i.e., ASD and TD) within each of the 10 strata. Note that, in the present work, the reason behind stratifying the subjects in our dataset based on their estimated propensity scores was different from that used in observational studies, where stratification can be employed for example to estimate stratum-specific effects. Stratification was here repurposed as a tool for the sole aim of training and testing predictive models on sets of subjects whose similarities and differences we could control in our data-analysis experiments.

### Out-of-distribution prediction

To illustrate the role of diversity in single-subject prediction, we started by comparing distinct subject sampling schemes when forming the training and holdout sets. In the *contiguous* scheme, we only selected adjacent strata for training and the remaining strata were “kept in a vault” for later testing of the generalizability in classifying ASD from TD. In the *diverse* scheme, the training set was composed of (at least one) non-contiguous strata. As a baseline, we furthermore used a *random* scheme where training data was drawn by taking into account the class proportions but irrespective of their propensity scores or strata. The *random* scheme amounted to a conventional cross-validation strategy.

This analysis was repeated for different sizes of the training set, ranging from 2 to 8 strata. In the *random* scheme, we hence pulled 20%-80% of the total subjects in our dataset for training of the predictive model. For each number of strata (or subset of subjects in case of *random* sampling) used for training, we performed 20 draws and computed the average of the ensuing model performances. In the *diverse* scheme, we did not consider all possible combinations of training/test strata, but rather focused on the combinations that provided the most diverse strata for training (i.e., those with the largest mean absolute difference in propensity scores). In the *contiguous* scheme, the number of possible draws was less than 20 (e.g., 9 maximum draws when using 2 contiguous strata for training).

To ensure a consistent number of draws, additional draws with similar size for the training set (e.g., 20% of pairs when using 2 strata for training) were sampled by considering paired subjects in consecutive order (according to their propensity scores) for training and the remaining for holdout. These sampling schemes allowed us to compare out-of-distribution (i.e., subjects with unseen propensity scores) performance when we learned from data that spanned the wider space parameterized by propensity scores (i.e., *diverse*), focused on a narrow subspace (i.e., *contiguous*), or completely ignored this kind of information in predictive modeling (i.e., *random*). Moreover, we also assessed performance within distribution. That is, based solely on the training strata, we used a 10-fold crossvalidation to report the performance when our model was learned and tested using data with similar propensity scores.

### Assessing the relationship of performance with diversity

The previous analyses offered an initial depiction of the implications of subject diversity on out-of-distribution performance. Yet, the analysis only considered 20 draws of training/test strata. In such case the *diverse* scheme may have been dominated by the strata at both extremes of the diversity spectrum, i.e., those with the lowest and highest propensity scores. This may have biased the results in terms of performance and also in terms of the stability of the predictive patterns, since the extreme strata were likely to be part of these 20 draws. Furthermore, diversity was experimentally controlled with a focus on the training set, rather than the holdout subjects. In another set of analyses, we examined the relationship between model performance and diversity for each draw of training/test strata. To this end, we used 5 strata for training (and the remaining 5 for holdout) and considered all possible combinations of subject strata to form the training set (i.e., 252 different draws, picking 5 out of 10 strata to form the training set).

Within-distribution performance was reported based on the average performance using a 10-fold CV strategy, whereas out-of-distribution performance was reported separately for each holdout stratum. For assessment of within-distribution performance, diversity was computed as the average of all pairwise absolute differences in propensity scores:

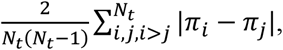

where *N_t_* is the total number of subjects in the training strata, and *π_i_*, denotes the propensity score of the *i*-th subject. For the purpose of out-of-distribution prediction, diversity denotes the mean absolute difference in propensity scores between all pairs of subjects in the training set and those in the holdout stratum:

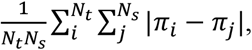

where *N_s_* is the number of subjects in the holdout stratum. This analysis allowed us to provide a clearer look into the impact of diversity on predictive modeling, by directly inspecting the relationship between subject diversity and predictive accuracy.

### Relation to common deconfounding practices

The covariates used in this study to investigate the impact of diversity in the classification of ASD may have a confounding effect in how the brain scanning measurements are used to predict diagnostic categories. In fact, numerous earlier studies have tried to account for nuisance sources by deconfounding the imaging-derived variables in what is often thought as a data-cleaning step (Alfaro-Almagro et al., 2020; Bernardino et al., 2020; Bzdok et al., 2020; Snoek et al., 2019). For instance, considering the sex of the subjects, we could find that males are more prone to be classified as ASD than females. This can be seen, for example, in the distribution of sex across the different strata in the HBN dataset (see **Figure 1A**). We noted that females occurred in the strata with the lowest propensity scores, while only males were in the highest propensity score strata. Regarding scanning site as a dimension of diversity, the variance of the brain-imaging-derived variables may differ from one site to another. This could lead to a biased estimation of the underlying ASD-related pattern in the brain since the model may have learned differences in variance rather than the true putative pattern (Dinga et al., 2020; Görgen et al., 2018). In such scenarios, it is important to seek to remove these sources of confounding.

To shed light on these circumstances, we explored: *i)* a conventional linear regression model and *ii)* ComBat (Johnson et al., 2007) for the purpose of variable deconfounding before the actual predictive model of interest. The first method is a standard approach to control for confounds in neuroimaging. It estimates the effect of confounds using a linear regression model to then remove their contribution to the data. The residualized functional connectivity strengths or regional morphological measurements then served as input variables of interest fed into the quantitative model of interest, such as a machine learning model (Dukart et al., 2011; Snoek et al., 2019). For categorical confounds such as the imaging site, this amounts to centering the data within each site, without considering the potential inter-site differences in variance (Dinga et al., 2020). To deal with the latter point, we also carried out ComBat: a recently proposed approach for better correcting inter-site effects (Fortin et al., 2017; Johnson et al., 2007). Briefly, ComBat involves Bayesian hierarchical regression modeling to estimate and remove site additive (i.e., shift in mean) and structured-noise effects (i.e., heteroscedasticity) from the brain-imaging features. ComBat has been previously applied to multiple neuroimaging data modalities including cortical thickness (Fortin et al., 2018; Larivière et al., 2020), resting-state functional connectivity (Yamashita et al., 2019; Yu et al., 2018), and diffusion tensor imaging (Fortin et al., 2017; Hatton et al., 2020).

We performed deconfounding using all subjects in our dataset, prior to matching and running our analysis. Note that in alternative settings, to prevent data leakage, only the training data should be used to build the deconfounding model, without considering the test data (Chyzhyk et al., 2018). However, the main reason behind our choice was that deconfounding in data points matched based on the same covariates may have little impact on the results (Linn et al., 2016).

### Dissecting the impact of cohort diversity

To provide a richer picture into how diversity affects disease classification, we next used confusion matrices to tease apart the cases in which our predictive models succeeded or failed. Moreover, prediction success may be contingent on the different covariates used to build the propensity scores. Here, we laid out the implications of diversity by taking into consideration the covariates of the holdout subjects. Given a particular holdout stratum, prediction performance was assessed for subjects from each of the different scanning sites. The analogous procedure was carried out to study sex. We also inspected the changes in performance that were attributable to age differences between the subjects in the training set and each holdout stratum.

Finally, we analyzed the degree to which diversity affects the stability of the extracted predictive patterns. To this end, we performed the analysis at both the regional (i.e., 100 target regions) and network levels (i.e., 7 target canonical networks) as defined by the widely used Schaefer-Yeo atlas (Schaefer et al., 2018; Yeo et al., 2011). First, we extracted the coefficients (i.e., weight vectors) from our 252 predictive model instances (corresponding to all possible combinations of picking 5 out of 10 strata to form the training set). These model coefficients were then assigned to 5 different groups according to the diversity of the training set used to learn the models (i.e., the average of all pairwise absolute differences in propensity scores of the subjects in the training set). We tested for significant coefficient differences between groups using classical one-way analysis of variance (ANOVA) with 5 levels (i.e., 5 groups: from very low to very high diversity, no continuous input variables). At the region level, we ended up with 252 coefficients corresponding to our collection of models. Differences in coefficient estimates were analyzed on a region-by-region basis using separate ANOVAs. This approach allowed identifying the cortical regions whose variance in predictive contributions (across the 252 previously obtained predictive models) could be partitioned according to the diversity factor levels in a statistically defensible fashion. Note that in the case of functional connectivity, we used model coefficients for each node (average of the regression coefficients across all edges of a node), and the threshold-free cluster enhancement approach was used with 1000 permutations to correct for multiple comparisons (Smith & Nichols, 2009). At the network level, we analyzed the relationships between model coefficients and diversity by computing Pearson’s correlation coefficient between the network-aggregated coefficients of predictive models and the diversity of the training subjects (i.e., the average of all the pairwise absolute differences in propensity scores of the training observations). These systematic assessments were aimed to uncover which parts of the brain are most at stake when predictive models from heterogeneous subject observations are to be interpreted by the neuroscientific investigator.

## Results

We investigated how diversity plays out in classification of neuropsychiatric disorders using structural and functional brain-imaging profiles constructed from the Autism Brain Imaging Data Exchange (Di Martino et al., 2014, 2017) and the Healthy Brain Network dataset (Alexander et al., 2017). Specific site inclusion criteria and rigorous data quality control resulted in a total of 297 subjects (151/146 ASD/TD) from 4 different acquisition sites in ABIDE and 206 subjects (104/102 ASD/TD) from 3 sites in HBN (**Tables S1** and **S2**). Our image processing strategy involved the mapping of functional signals to cortical surfaces as well as surface-based alignment. Functional connectivity matrices were calculated at a single-subject level based on a widely used parcellation atlas with 100 cortical regions (Schaefer et al., 2018).

Prior to predictive modeling, ASD and TD subjects were matched and grouped into strata with homogeneous backgrounds based on their propensity scores (see **Figure 1**). This step yielded a total of 290 subjects (145 subjects per group) in ABIDE and 126 in HBN. In ABIDE, a very small number of subjects was discarded from subsequent analyses after matching by age, sex and acquisition site. However, with HBN around one third of the original subjects was excluded because of the modest overlap between the distributions of estimated propensity scores between TD and ASD groups. In **Figure 1A**, we can see that in HBN the distribution of TD subjects was skewed to the right while for ASD it was skewed to the left. In HBN, we further explored the role of diversity in the subject classification of ADHD vs TD and ANX vs TD. After matching, we submitted a total of 180 (90 ADHD/TD) and 188 (94 ANX/TD) subjects to each classification analysis, respectively. Further information about the datasets, image processing, matching and classification settings is provided in the *Methods* section.

### Benchmarking out-of-distribution prediction performance

Results in the classification of ASD versus TD subjects in both ABIDE and HBN datasets are shown in **Figure 2**, with performance reported in terms of area under the receiver operating characteristic curve (hereafter, AUC) and F1 score. In the ABIDE cohort, when we trained and tested on data with similar distributions (first column), the *contiguous* scheme reached better prediction performance than the *diverse* scheme, with the *random* scheme lying in between. This trend remained stable as we increased the number of strata used for training. However, we can observe in the holdout dataset (second column), that the difference in prediction accuracy showed an inverted trend. The predictive model trained on the contiguous strata showed poor generalization abilities. We note a clear decay in classification performance as we increased the number of strata used for training, which may suggest that the model was overfitting to subjects that share similar demographic status (e.g., age and sex), and hence may have been limited to generalize to more diverse data. On the other hand, the *diverse* and *random* schemes performed better as we grew the size of the training set. In the HBN dataset, we observed the opposite trends with the *diverse* analysis scenario, which showed higher performance within-distribution and poor generalizability.

**Figure 2.**
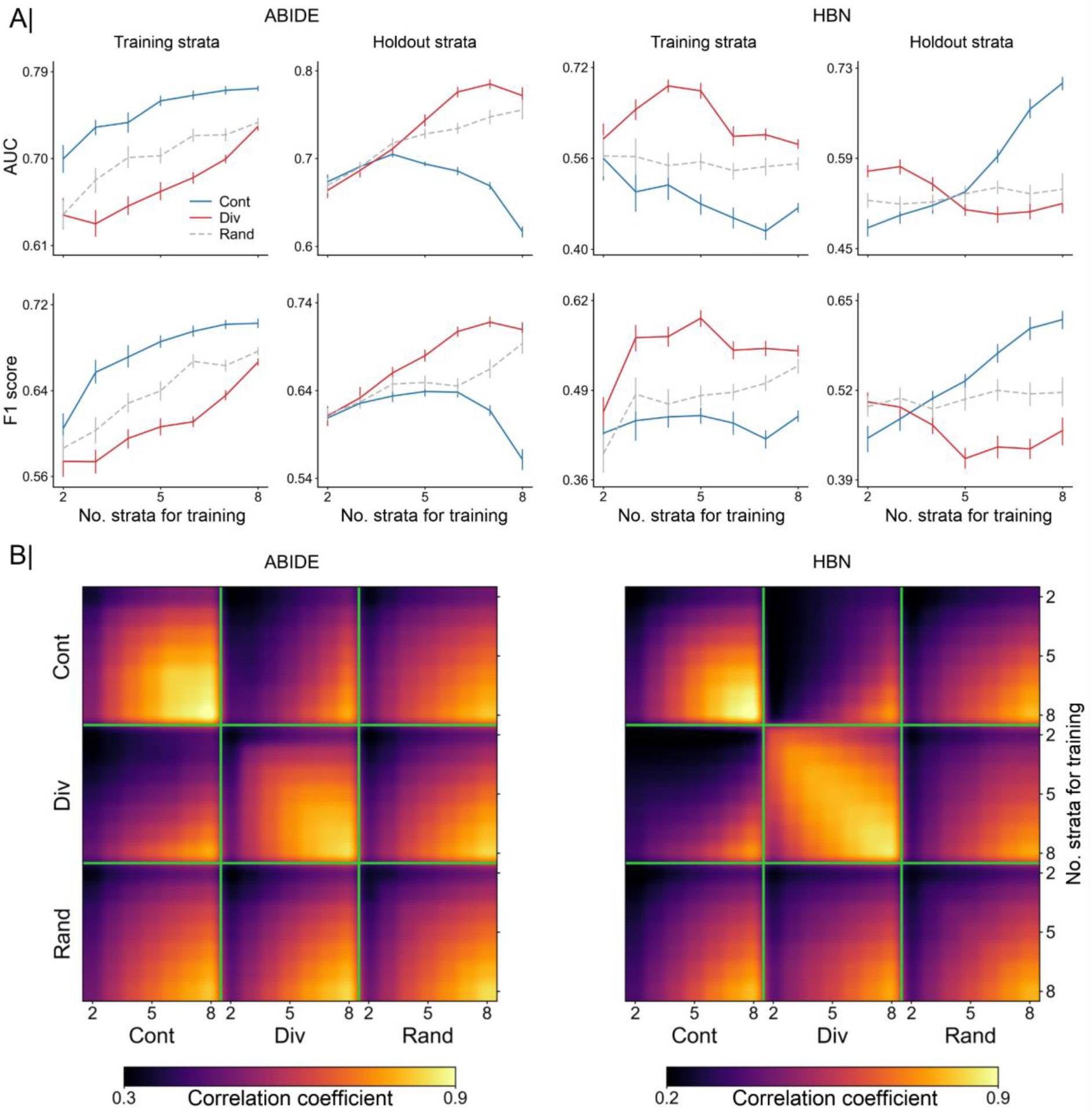
Accuracy of out-of-distribution prediction and consistency of extracted predictive patterns. A) Comparison of model accuracy based on contiguous (Cont) and diverse (Div) training sets in the classification of autism. Prediction performance is reported as area under the curve (AUC, top) and F1 score (bottom) in two separate cohorts: ABIDE (left) and HBN (right). Accuracy is assessed using different training sets sized from 2 to 8 combined strata. For each cohort, the first column indicates the prediction accuracy using a 10-fold crossvalidation strategy based solely on subjects from the training strata. Folds were randomly sampled, without considering the propensity scores of the subjects. The second column displays the performance in the holdout strata, which contains the remaining subjects (from untouched strata). As a baseline for comparison, we used an additional sampling scheme (Rand): subjects for the training set were randomly chosen regardless of their propensity scores. B) Consistency of model coefficients was quantified by Pearson’s correlation coefficient across two given models in the contiguous, diverse, and random data scenarios. For each of these sampling schemes, consistency is shown for different numbers of combined strata used for predictive model training (from 2 to 8 combined strata), delineated by the green segments. Building models based on diverse strata of subjects entailed considerable differences in predictive patterns from those learned based on models with similar subjects (i..e, contiguous).

In the context of these diverging results between ABIDE and HBN, we note the number of females in each dataset (ABIDE 5/290, HBN: 63/126). As shown in **Figure S1**, where we display the distribution of sex across the strata, in HBN all females were confined to the first 6 strata (those with the lowest propensity scores). In contrast, in strata 7 to 10, there were only males. This circumstance may explain the drop in out-of-distribution performance in the *diverse* scenario because of the imbalanced sex covariate. Nonetheless, in both datasets we observed a strong impact of diversity in model performance. This finding indicates that within-distribution performance is not in all cases an accurate proxy for out-of-distribution performance when there is a shift in the distribution of the covariates.

Such tendency received further support from our results in classifying ASD vs TD using cortical thickness in the ABIDE dataset, and the other two neuropsychiatric disorders (i.e., ADHD vs TD and ANX vs TD) from the HBN dataset based on functional connectivity (see **Figure S4**). With cortical thickness, we found similar performance trends as the ones obtained with functional connectivity in ABIDE, with the *contiguous* scheme outperforming the *diverse* scheme within-distribution but showing lower accuracy in the holdout subjects. The same performance pattern was found in ANX. On the other hand, ADHD showed similar trends to the ones found in the classification of ASD in HBN. Model accuracy in the classification of ADHD, however, was very close to 0.5, which may suggest that there may not be enough signal in the data.

We also analyzed the relationship between the functional connectivity patterns (i.e., weights) learned by the models based on the two different sampling schemes. More specifically, we calculated the Pearson’s correlation coefficient to quantify the consistency of model coefficients that we obtained in the scenarios with *contiguous*, *diverse* and *random* sampling schemes. Results are shown in **Figure 2B** for different sizes of the training set (from 2 to 8 strata for model training) in both ABIDE and HBN datasets for the classification of ASD. With the exception of 2 strata, where the model may not have enough data to extract a robust pattern, the models were able to identify robust patterns (correlation coefficient r>0.7) in both datasets with the *diverse* sampling. With the *contiguous* and *random* schemes, the extracted patterns showed high consistency only when large numbers of strata were used for training, although in ABIDE the *contiguous* scheme was able to extract stable patterns. This is mainly due to the overlap of the training data, which may relate to our observation of strong pattern stability comparing the sampling schemes when using a large number of strata for training (e.g., 8 strata). Overall, however, the patterns identified by the *contiguous* and *diverse* schemes showed very low correlations, which highlights the important role of diversity in obtaining robust and reproducible biomarkers. This was also illustrated by the low pattern correlations of the *random* scheme, where training subjects were chosen without considering the criterion of diversity. **Figure S5** shows the consistency of patterns obtained in the classification of ASD using cortical thickness in the ABIDE dataset, and ADHD and ANX from HBN based on functional connectivity. In accordance with the previous results obtained in the classification of ASD based on functional connectivity, we found a low consistency between the patterns extracted from contiguous and those from diverse strata.

### Charting the tension between prediction accuracy and subject diversity

**Figure 3** displays the relationship between model performance and diversity in both ABIDE and HBN datasets in the classification of ASD from functional connectivity. For each cohort, the first column shows within-distribution performance based on a 10-fold CV strategy using only the training set, and the second column shows performance in the holdout data. Results were obtained using 5 strata to build our predictive models (and the remaining 5 for holdout). Performance is reported in terms of AUC and F1 score. With both performance metrics, we can observe a decline in performance in ABIDE as diversity increased in both within (AUC: r=-0.630, F1 score: r=-0.568) and out-of-distribution (AUC: r=-0.671, F1 score: r=-0.579) predictions. In the HBN cohort, we found the opposite relationship, with performance being negatively correlated with diversity when assessing both within (AUC: r=0.427, F1 score: r=0.413) and out-of-distribution (AUC: r=0.414, F1 score: r=0.397) predictions. In both datasets we found strong associations between performance and diversity, although with different signs. In light of the diverging results observed in the two cohorts and the negligible number of females in ABIDE, we further inspected the role of diversity in the HBN dataset when considering males and females separately. Diversity was thus only defined by age and scanning site. As illustrated in **Figure S2**, performance (both within and out-of-distribution) decayed with increasing diversity in both female and male subsets, showing therefore similar trends to that found in the ABIDE cohort. Moreover, **Figure S4** shows the relationship between performance and diversity in the classification of ASD in ABIDE based on cortical thickness, and ADHD and ANX in HBN using functional connectivity. In ADHD, we can observe a positive correlation of performance with diversity. Yet, it was very weak in holdout subjects. For the two other cases (i.e., ASD vs TD using cortical thickness and ANX vs TD using functional connectivity), we found negative correlations, with performance decreasing the farther the held-out stratum was from the strata used for training.

**Figure 3.**
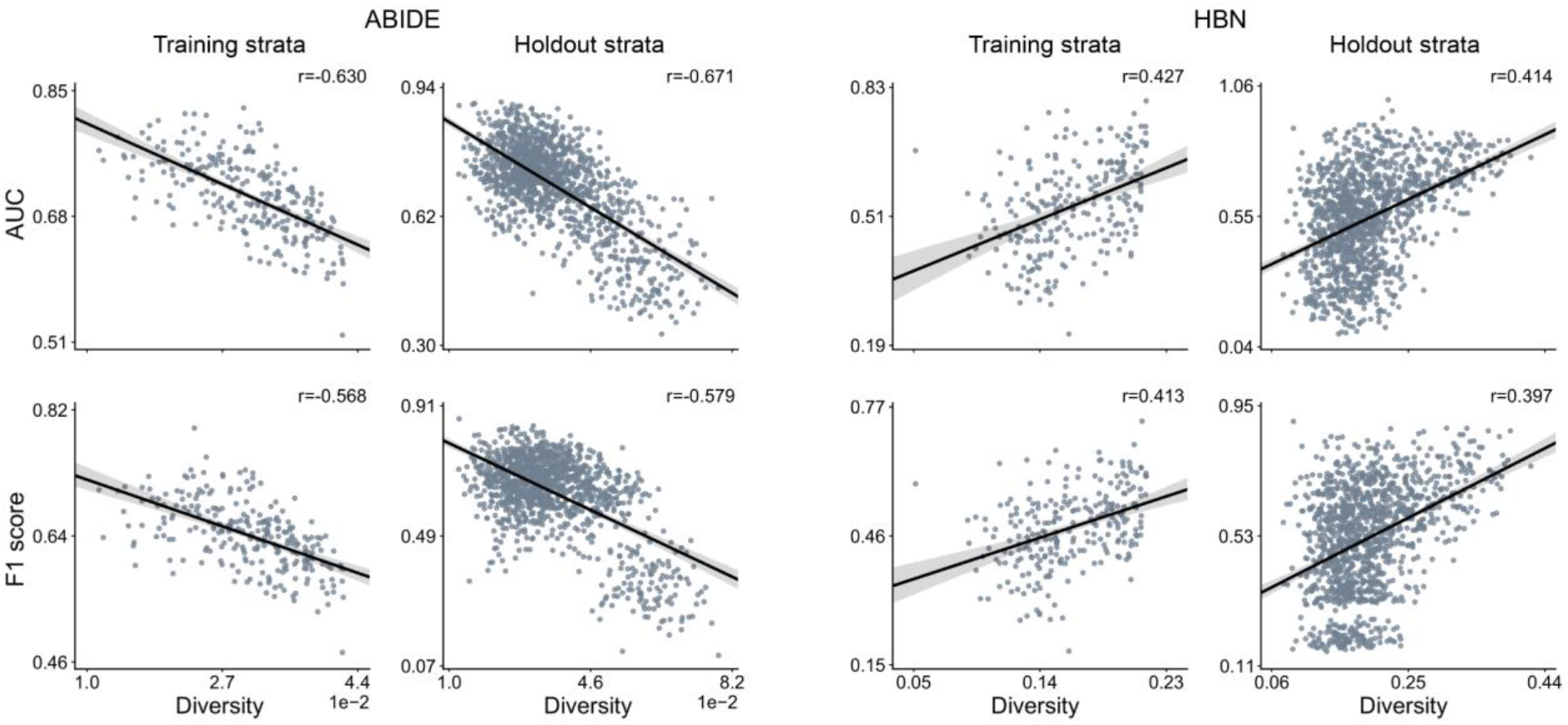
Subject diversity is a major determinant for the classification accuracy of predictive models. Results are based on all possible combinations of 5 out of 10 strata for training and the remaining 5 strata as holdout. Prediction accuracy based on area under the curve (AUC, top) and F1 score (bottom) in two different cohorts: ABIDE (left) and HBN (right). For each cohort, the first column indicates the predictive model performance using a 10-fold cross-validation strategy based solely on the training set, where diversity is computed as the average of all pairwise absolute differences in propensity scores. The second column displays the performance for each single stratum in the holdout strata. Diversity denotes the mean absolute difference in propensity scores between the subjects of the training set and those in the held-out strata with unseen subjects. The strength of the association between performance and diversity is reported with Pearson’s correlation coefficient (r). Our empirical results show a strong relationship between predictive performance and diversity, although different correlation directions were found in ABIDE and HBN cohorts.

### Exploring the role of deconfounding practices

So far, we have shown that diversity had a considerable impact on model performance, but we did this based on the original raw functional connectivity data. In this section, our aim is to investigate if deconfounding can help us remove, at least to some extent, the impact of diversity on performance. To do so, we used 2 different deconfounding approaches. The first deconfounding approach is the traditional linear-regression-based approach that removes the contribution of the confounds from the raw brain-imaging data. For the second deconfounding approach, we used ComBat to deal with site effects. Similar to the computation of the propensity scores, we performed deconfounding prior to our analyses, based on all the available subjects instead of using only the training data. The relationship of performance with diversity after deconfounding is reported using AUC in **Figure 4** and F1 score in **Figure S3** in the supplementary materials. We refer to the absence of deconfounding as *raw* data (first row). In ABIDE, the correlations of AUC and diversity in the holdout dataset when using linear-regression-based (r=-0.539) and ComBat (r=-0.599) deconfounding approaches were slightly lower than the correlation obtained when using the raw data (r=-0.671). In HBN, we also found small differences with the linear-regression-based (r=0.327) and ComBat (r=0.392) approaches than when using the raw data (r=0.414). Similar results were obtained with F1 score. It is worth mentioning here that although there were slight differences, we cannot expect to find major changes because our data was already matched. Nonetheless, these results further emphasized the importance of taking diversity into consideration when using cross-validation strategies to assess the performance of our predictive models.

**Figure 4.**
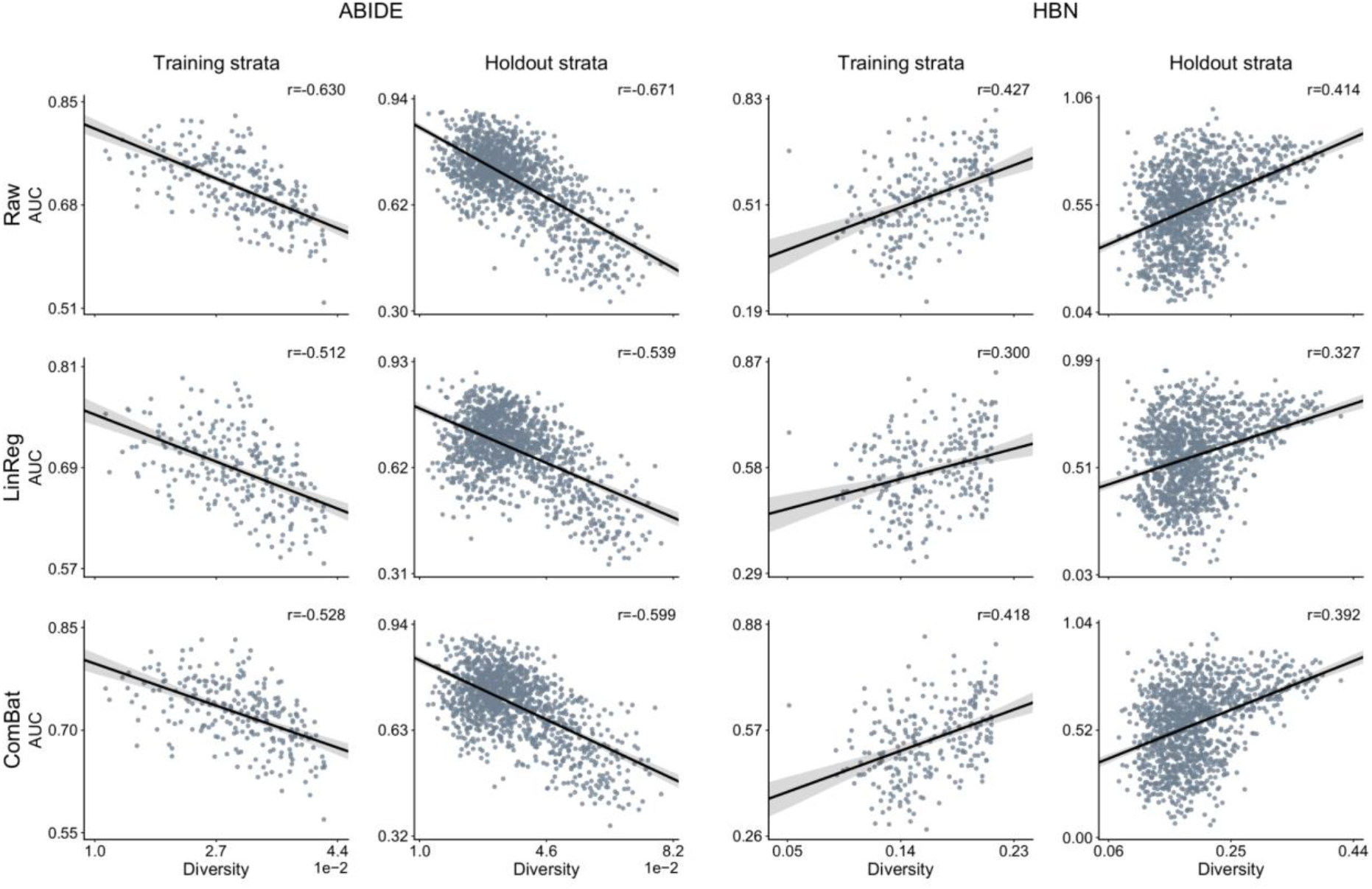
Established and new deconfounding strategies appear insufficient to counteract escalating population diversity. A means to quantify the behavior of predictive models is comparing its predictions on training and testing information. Performance is measured by area under the curve (AUC) metric on the original brain-imaging data (*Raw*, top), and after carrying out deconfounding steps on the brain feature prior to the pattern classification pipeline using standard linear-regression-based deconfounding (*LinReg*, middle) and recently proposed ComBat (bottom) approaches. Compared with the conventional nuisance removal using regression residuals, ComBat is a more advanced hierarchical regression to control for site differences. Results are reported for two different clinical cohorts: ABIDE (left) and HBN (right). For each cohort, the first column shows the model prediction performance using a 10-fold cross-validation (CV) strategy. Diversity is computed as the average of pairwise absolute differences in propensity scores between all subjects. Each dot is a cross-validated accuracy. The second column, for each cohort, displays the performance in the holdout subjects. Instead of reporting performance on all the subjects in the holdout data, here performance is assessed independently for subjects in each single stratum from the holdout data. Thus, diversity denotes the mean absolute difference in propensity scores between the subjects in the training set and those in the held-out stratum. Both ComBat and linear-regression-based deconfounding failed to mitigate the impact of diversity on prediction accuracy.

### Dissecting the impact of diversity in prediction accuracy

To better understand the impact of diversity in performance, we inspected the out-of-distribution performance of our predictive models using confusion matrices. First, the holdout strata were ranked and divided into six chunks according to their diversity (with respect to the training strata) to then generate the respective average confusion matrices. **Figure 5A** shows out-of-distribution confusion matrices in the classification of ASD in both ABIDE and HBN datasets. In ABIDE, with low diversity (e.g., the first confusion matrices) we see that the predictive models achieved good discriminative power, with a proportion of true positives (TP) and true negatives (TN) of 0.33 and 0.35, respectively. But as diversity increased, the model tended to classify most subjects as TD. In the confusion matrix corresponding to the highest diversity, the proportion of TN was 0.37 and that of false positives (FP) was 0.38. In the HBN dataset, the impact of diversity was reflected in the opposite direction, starting with low performances (e.g., proportion of TP and TN in the first confusion matrix of 0.22 and 0.25) to then achieve better predictions as diversity increased (e.g., proportion of TP and TN of 0.33 and 0.39 in the last confusion matrix).

**Figure 5.**
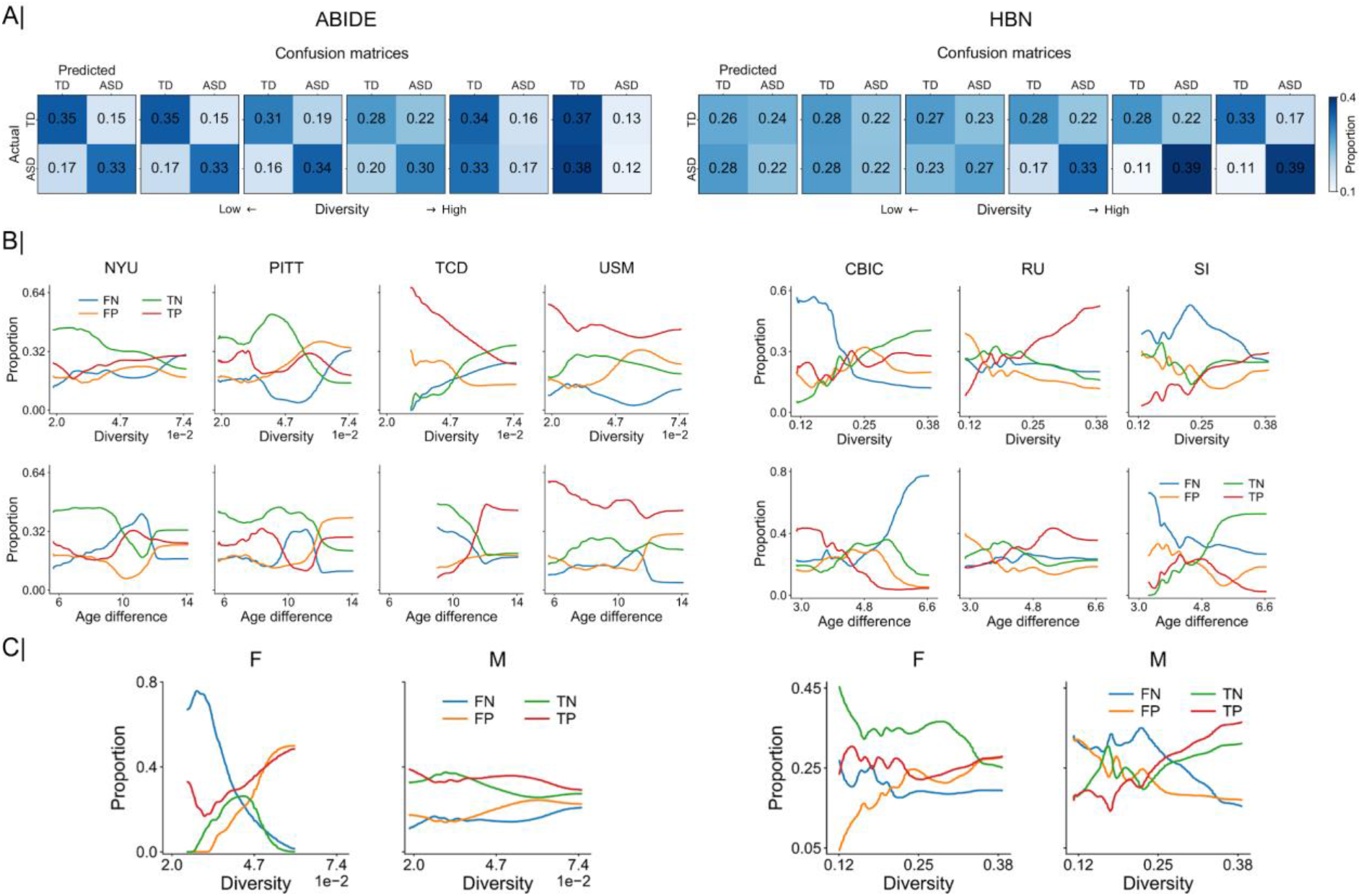
Break-down of out-of-distribution predictions reveals unstable model behavior according to diversity subdimensions: acquisition sites, subject age and sex. A) Error matrices that summarize the relationship of diversity with success and failure rate of the predictive models. These matrices are computed across sets of diagnostic classifications sorted in ascending order by diversity. B) Relationship of out-of-distribution performance with diversity and age for each scanning site in ABIDE (left; NYU, PITT, TCD and USM) and HBN (right; CBIC, RU and SI). Performance is shown in terms of proportion of false negatives (FN), false positives (FP), true negatives (TN), and true positives (TP) cases of model-based classification of clinical diagnoses. C) Relationship of out-of-distribution performance with diversity for female (F) and male (M) subjects. Predictive models show unstable behavior across each diversity dimension, underscoring the inconsistent results across the different sites and between females and males. Abbreviations: NYU, New York University Langone Medical Center; PITT, University of Pittsburgh, School of Medicine; TCD, Trinity Centre for Health Sciences, Trinity College Dublin; USM, University of Utah, School of Medicine; SI, Staten Island; RU, Rutgers University Brain Imaging Center; CBIC, CitiGroup Corcell Brain Imaging Center.

We also examined the impact of diversity for each of the covariates that contributed to its computation (i.e., age, sex and site). As shown in **Figure 5B**, the impact of diversity in classification performance (in terms of true/false positives/negatives) varied from one scanning site to another. Considering, for example, the proportion of TN in ABIDE, we can see that it decreased with diversity in most sites except in TCD, where it did increase. Similar results could be found in HBN, where the proportion of TN increased with diversity in one single scanning site (i.e., CBIC) but decayed in the remaining sites. This indicates that diversity impacts the performance in each site differently, which might account for the divergent results observed in ABIDE and HBN. When plotting the same performance results against age, we found different trends. This occurred because the distribution of age across the strata was not monotonically increasing with diversity, as shown in **Figure S1**. Nonetheless, in HBN for instance, we can see that the proportion of false negatives (FN) increased with age in CBIC while it decreased in the rest of the scanning sites. Finally, when using sex (see **Figure 5C**), we found a clear decrease in the proportion of FN in females while a slight improvement was observed in males in the ABIDE dataset. Note however that there was a substantially reduced number of females in ABIDE compared to HBN. In HBN, we see that the proportion of FP, for example, increased in females and decreased in males. Since we are only interested in the overall trends, the results in **Figures 5B** and **5C** were smoothed using a Gaussian-weighted moving average for visualization purposes.

### Quantifying the impact of diversity on pattern consistency

**Figure 6A** displays the coefficients extracted by the predictive models with increasing diversity in both ABIDE and HBN cohorts. Model coefficients were z-scored, grouped into five chunks and averaged according to diversity, with positive and negative coefficients shown separately. In both datasets, we can observe that these coefficients underwent considerable drifts in the face of diversity. For example, the coefficients of both the medial frontal and posterior cingulate cortices changed considerably with diversity, going from positive to negative in ABIDE and vice versa in HBN. Moreover, **Figure 6B** illustrates how consistency changed between patterns extracted from subjects in the training set with different diversity. We can see that the extracted patterns gradually diverged as diversity in the training data increased. This is shown in both datasets, but more markedly in HBN. Similar results can be seen in **Figure S6** in the classification of ASD using cortical thickness and ADHD and ANX based on functional connectivity. The regions whose coefficients underwent significant changes with diversity are shown in **Figure 7A** and **7B** for ABIDE and HBN, respectively. Although this is not directly related to the discriminative importance of such regions, it does show that the patterns extracted from homogeneous observations differ from those obtained when training with more heterogeneous observations. At the network level, we found strong associations of network coefficients with diversity in the default mode (consistency of predictive patterns: r=-0.286/0.161 in ABIDE/HBN), frontoparietal (r=0.413/-0.291), limbic (r=0.373/0.110), and dorsal (r=0.137/-0.247) and ventral (r=0.298/0.330) attention networks. There does not seem to be a strong effect of diversity in the model coefficients corresponding to the somatomotor (r=-0.112/-0.099) and visual (r=0.046/-0.108) networks.

**Figure 6.**
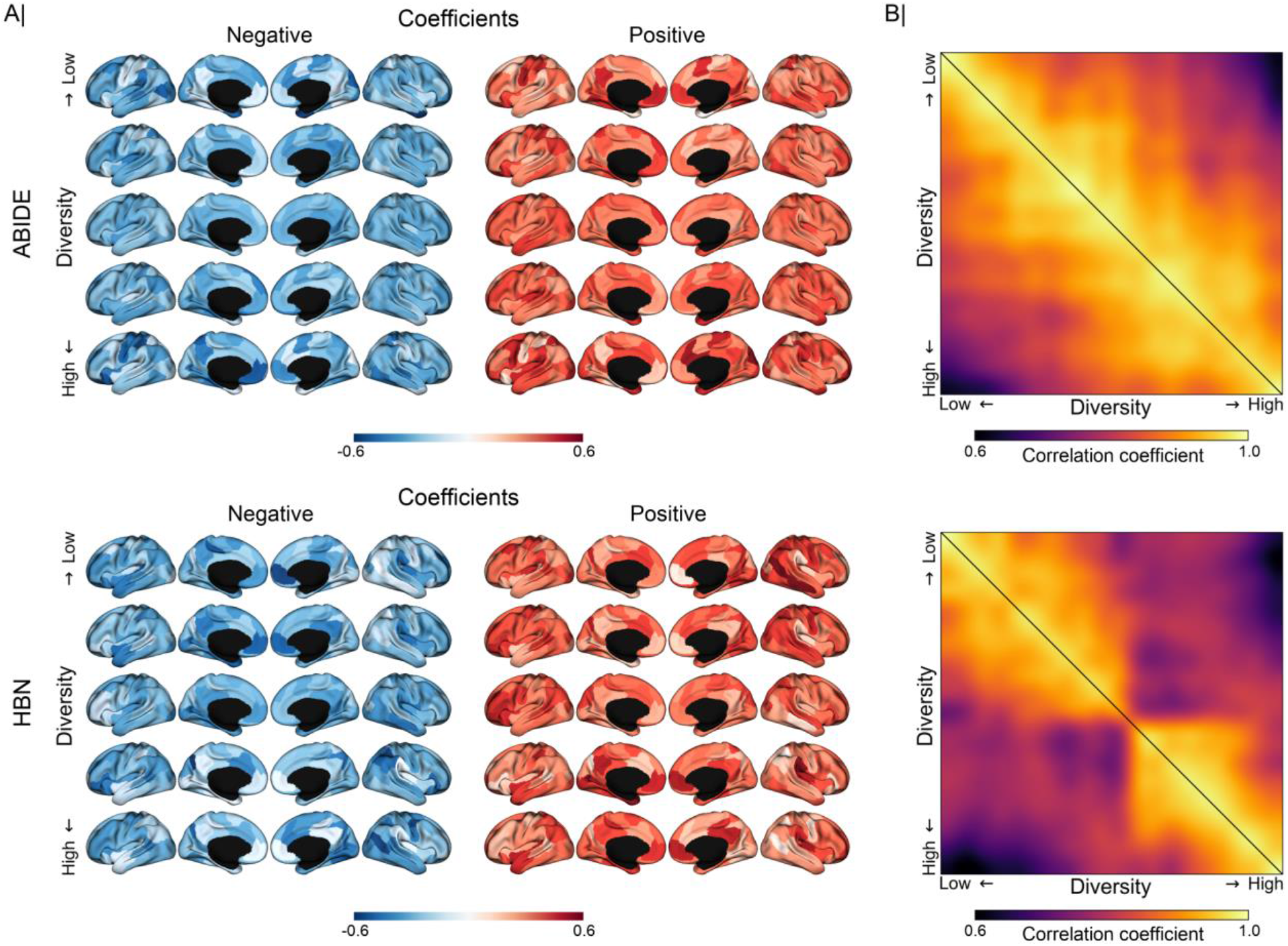
Varying subject diversity is detrimental for consistency of model-derived predictive patterns. Results are based on all possible combinations of 5 out of 10 strata for training and the remaining 5 strata as holdout. A) Drifts in model coefficients when estimated repeatedly (rows) with increasing population stratification, separately in ABIDE (top) and HBN (bottom) cohorts. Model coefficients were ranked according to diversity and grouped into five chunks. Coefficients averaged within each chunk are displayed with increasing diversity. Positive and negative coefficients are shown in separate brain renderings for visibility. For each node, positive/negative coefficients were computed by averaging the edges with only positive/negative coefficients. B) From top to bottom, changes in predictive model coefficients with increasing diversity in ABIDE and HBN cohorts. Consistency of model coefficients, in terms of Pearson’s correlation, is obtained for each possible combination of training subjects (5 strata combined for training), where diversity is computed as the mean absolute difference in the propensity scores of the training subjects. Each entry in the correlation matrices corresponds to the correlation between the coefficients obtained by two predictive models trained on different combinations of training strata. Model coefficients were sorted according to the diversity of their corresponding training observations (arranged from low to high). Each matrix shows correlations based on the model coefficients learned when using the raw data (lower triangular part) and the ComBat-deconfounded data (upper part). Our results show that the consistency of model-derived predictive patterns decays with increasing diversity of the training set, even under deconfounding.

**Figure 7.**
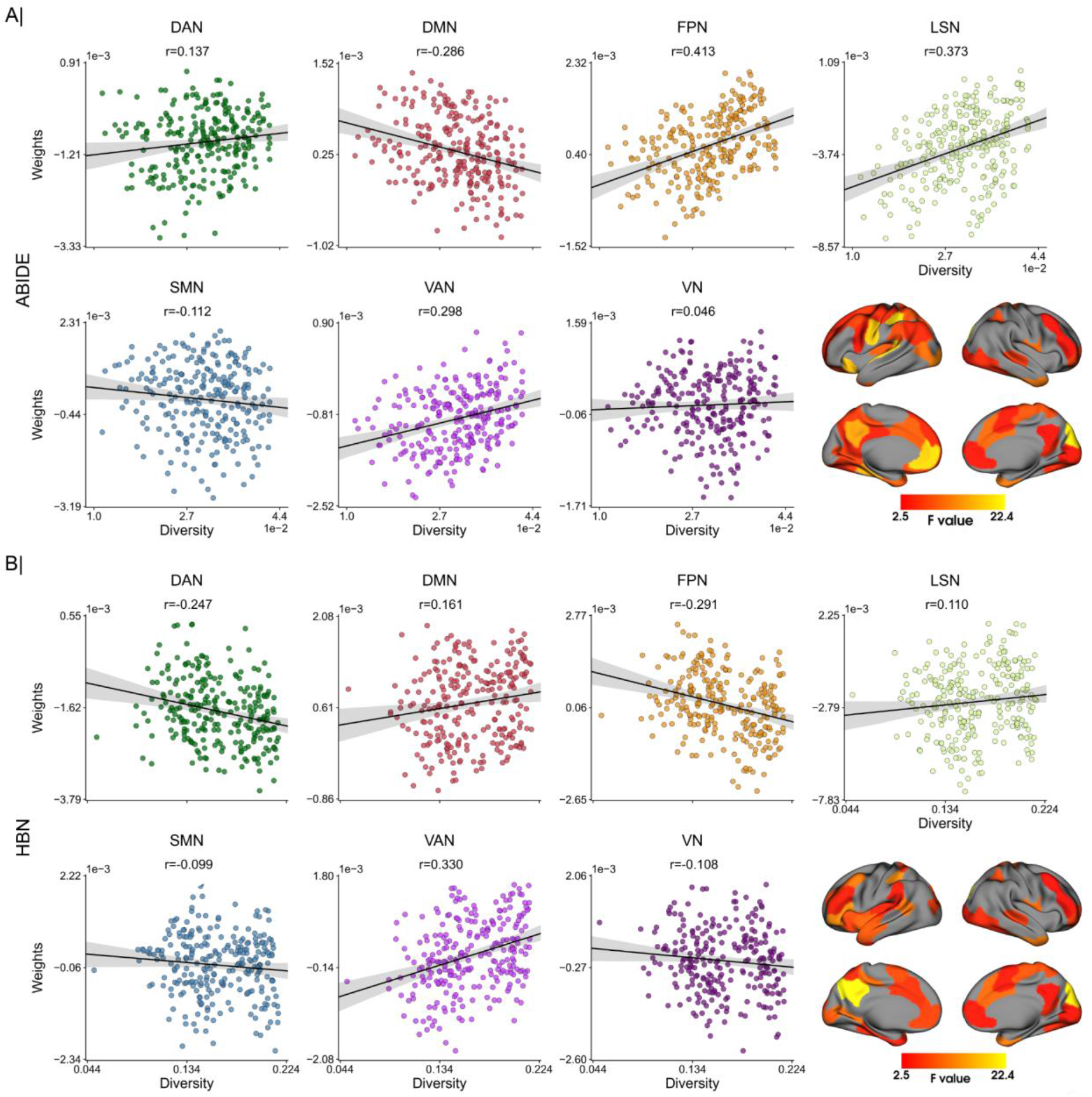
Anatomical hot spots where predictive rules risk to become brittle in the face of diversity. ABIDE cohort: Relationship between network-aggregated coefficients of predictive model and level of population diversity. Model coefficients are averaged for each intrinsic connectivity network. B) HBN cohort: Relationship of network-aggregated model coefficients with diversity. Brain renderings in A and B expose regions whose coefficients show a significant association with diversity. Consistent across both clinical cohorts, predictive model coefficients in regions of the highly associative default-mode network showed a privileged relation to escalating population stratification. Abbreviations: DAN, dorsal attention network; DMN, default mode network; FPN, frontoparietal network; LSN, limbic system network; SMN, somatomotor network; VAN, ventral attention network; VN, visual network.

## Discussion

In our quest towards realizing single-patient prediction in real-world settings, MRI-based machine learning approaches seek to provide accurate and replicable biomarkers. Due to the recent rise of large-scale brain scanning collections, analytical tools are now urgently needed to account for potential *distributional shifts* as a consequence of increasing diversity of the subject cohorts. In multi-site neuroimaging studies, the generalization power of the predictive models is more likely to be affected by several sources of population stratification. Some previous research studied their impact on prediction accuracy and biomarker robustness (Bzdok & Meyer-Lindenberg, 2018; Dinga et al., 2020; Lanka et al., 2019). Yet, such analyses typically focused on a single source of subject heterogeneity. Our study sought to accommodate cohort diversity coming from multiple sources simultaneously to examine *out-of-distribution generalization*. We conducted computational experiments that delineate how success and failure of brain-based machine-learning predictions are conditioned on sample diversity based on brain-imaging-derived profiles of structure and function in two different ASD cohorts. Our findings spotlight the higher-order association cortex, especially the default mode network, as the candidate substrate where subject diversity was linked to the biggest instability of extracted predictive signatures. Our results further show that this interdependence was not only specific to ASD and functional connectivity. Predictive signatures were also driven by subject diversity when studying structural features (i.e., cortical thickness), and functional connectivity in other neuropsychiatric disorders (i.e., ADHD and ANX).

For the purpose of the present work, we construed diversity in terms of three key covariates: age, sex and brain scan acquisition site. These covariates are well known to affect neuroimaging results (Alfaro-Almagro et al., 2020; Bzdok et al., 2020; Duncan & Northoff, 2013). Since ASD is a neurodevelopmental disorder with potential compensatory mechanisms that occur across the lifespan (Lord et al., 2015; Ullman & Pullman, 2015), learning an age-independent biomarker from the brain measurements remains a challenging task. From group-level contrast analyses, we know that the form and direction of the alterations found in ASD vary throughout development in both structural (Khundrakpam et al., 2017; Nunes et al., 2020; Zielinski et al., 2014) and functional (Kozhemiako et al., 2020; Nomi & Uddin, 2015) brain-imaging. Thus, the complex nature of these disease-related brain correlates hinders the generalizability of machine learning methods across age. These circumstances probably surface as lower classification accuracies that are achieved in subjects from different age groups (Lanka et al., 2019; Vigneshwaran et al., 2015). Brain manifestations related to ASD were also found to diverge between males and females (Alaerts et al., 2016; Lawrence et al., 2019; Retico et al., 2016). In fact, in ASD, there is a well-known prevalence disparity, with an approximate sex ratio of 4 males per female (Blumberg et al., 2013). This aetiological observation points to the idea that sex is an important contributor to the diagnosis. Moreover, higher prevalence of males is observed in ADHD (Ramtekkar et al., 2010), while ANX is more common in females (McLean et al., 2011).

Additionally, when using multi-site data, sites may be at odds with respect to other covariates such as age and sex. In our ABIDE cohort, there was only one site that contributed females. Hence, using data from different acquisition sites leads to potential heterogeneity attributable to sampling bias (i.e., demographics) and measurement bias including factors such as scanner type and imaging protocol (Yamashita et al., 2019). In a similar vein, several studies have reported declines in classification performance when using multiple sites (Colby et al., 2012; Lanka et al., 2019; Nielsen et al., 2013). By designing and executing a framework that jointly considered all these covariates simultaneously, we noticed a considerable drop in out-of-distribution performance in the classification of ASD in ABIDE, based on both functional connectivity and cortical thickness measurements, as well as in patients with ANX in HBN. In contrast, our predictive models achieved higher performance in out-of-distribution classification in patients with ASD and ADHD in HBN. The incongruent results in ASD classification performance between ABIDE and HBN may in part be explained by reduced size of the data in HBN after matching (126 subjects in HBN versus 290 in ABIDE). Other factors that might have contributed to this difference include the varying number of females in each dataset (with a considerably higher prevalence in HBN), as well as the different sample sizes available from each site.

In the HBN cohort, our subanalyses on individuals only from one sex showed a similar constellation of findings to those in ABIDE (in both female and male subsets of HBN). However, in the more challenging, yet more realistic setting of multi-site datasets, classification accuracy within each site has been shown previously to be highly variable (Spera et al., 2019), and sites that account for a large portion of the observations may therefore drive prediction performance. Inter-site differences also became apparent when we further dissected the role of diversity on performance, where the relationship of performance and diversity varied from one site to another as well as between males and females in both ABIDE and HBN. Overall, we note a strong interplay between prediction accuracy and diversity in the population under study, although inconsistent across its different dimensions. These insights from two of the largest autism brain-imaging cohorts underscore the need to take into account multiple dimensions of subject stratification that may potentially contribute to diversity when testing the generalizability of a predictive model, rather than only considering one single factor at a time.

As a central neuroscientific finding from our investigation, the extracted predictive patterns have flagged a coherent set of brain substrates to be associated with subject diversity, which was consistent across ABIDE and HBN cohorts. The robust brain manifestations of diversity included especially regions of the default mode network, as well as the frontoparietal and attention networks. The maturational processes of these associative regions are thought to be more prolonged than those seen in primary sensory or motor brain circuits, which are established earlier in life (Gogtay et al., 2004; Wierenga et al., 2016). A more gradual maturation trajectory makes for a wider window for the action of environmental factors that may contribute to the increased functional variability of these neural network systems (Mueller et al., 2013). Moreover, differences in myelin content are also observed between these canonical networks, with the higher association cortex showing fewer myelinated axons than primary sensory and motor cortices. Less myelination might enable greater plasticity in association areas both during development and adulthood (Glasser & Van Essen, 2011; Timmler & Simons, 2019; Turner, 2019). Neural circuits with less myelinated regions have previously been shown to exhibit increased functional and structural variability (Karahan et al., 2021). Taken together, these factors might be related to, at least partly, the high susceptibility of our predictive models to population diversity in the default mode network, and other associative cortical regions.

More specifically, in ASD, these functional systems of the higher association cortex have frequently been reported to be disrupted in a long series of existing studies (Assaf et al., 2010; Di Martino et al., 2009; Farrant & Uddin, 2016; Fitzgerald et al., 2015; Hong et al., 2019; Just et al., 2012; Kennedy & Courchesne, 2008; Müller et al., 2011). Additionally, regions within these networks were found to be discriminative in machine-learning classification of ASD versus TD subjects (Abraham et al., 2017; Kernbach et al., 2018; Plitt et al., 2015). The high variability in the coefficients of these networks in relation with our diversity indicators is in line with the inconsistencies regarding hypo/hyper-connectivity findings in ASD. In the default mode, for example, the direction of the disease-related functional connectivity varies with the age of the cohort under study, shifting from hyper-connectivity in children to hypo-connectivity in adults, with both co-occurring in cohorts with broader age ranges (Padmanabhan et al., 2017). In addition, these networks are highly idiosyncratic in ASD, showing greater topographic variability among ASD individuals than TD controls (Benkarim et al., 2020; Dickie et al., 2018; Nunes et al., 2019). This increased idiosyncrasy may also contribute to this elusiveness of default network and other association regions to yield stable predictive signatures. Hence, our collective findings make apparent that diversity selectively impacts pattern stability especially for some of the highest integration centers of the human cerebral cortex, which we note to coincide with the functional systems that are, instead, routinely reported to be specific for patients with ASD.

The proof-of-principle analyses in this work were centered on age, sex and site because these indicators of diversity are among the frequently available in many biomedical datasets. However, there are several factors that could and should be considered in future research: verbal and non-verbal IQ, open vs. closed eyes during functional brain scanning, ethnicity/race, and comorbidity among others. These factors may play a pivotal role for classification accuracy and pattern stability of predictive models, from support vector classification to random forest algorithms to deep artificial neural networks. By benefitting from propensity scores, investigators can integrate the covariates or dimensions of population stratification into the analytical workflow to carefully monitor their implications for a statistical analysis of interest. Additionally, our study only considered the role of diversity in classification, but our analysis pipelines are also applicable to regression problems. In the latter, diversity can be computed based on generalized propensity scores (Bia & Mattei, 2008; Hirano & Imbens, 2004), which are an extension of propensity scores to work with continuous variables (Austin, 2018). Propensity scores could be used for instance in the prediction of symptom severity (e.g., ADOS) rather than diagnosis. Indeed, focusing on symptom severity instead of diagnosis may account for part of the heterogeneity observed in ASD (Moradi et al., 2017).

The generalization failure exposed by the present analyses highlights an important instance of *dataset shift* (Candela et al., 2009), a well-established concept in the machine learning community. Dataset shift occurs when the joint distribution of the test data differs from that of the training data. There are different types of dataset shift. In order to determine the type of shift we need to identify: *i)* the factors of the joint distribution that change between the training and testing stages of the predictive modeling workflow, and *ii)* the causal relationship between outcome variable and the brain features (Castro et al., 2020; Moreno-Torres et al., 2012). Thus, elucidating the causal structure of the variables at play in our problem would help identify susceptibility to shifts and build more robust predictive models (Subbaswamy & Saria, 2020). This realization highlights the importance of future work to exploit causal knowledge to better deal with potential distributional shifts. For example, incorporating causal knowledge has been shown to allow for better performance than the traditional linear-regression-based approach for deconfounding (Neto, 2020). In our case, repeating our analysis using different linear deconfounding routines only led to slight differences in the relationship between model performance and diversity. This observation underlines the limitations of the deconfounding approaches that the neuroimaging community is employing nowadays.

Ultimately, adjusting for the “right” confounding covariates is a causal matter at its heart. Our study goal has a direct relationship to the notion of *ignorability* from the causal inference literature - a form of independence that is sometimes also referred to as unconfoundedness (Pearl, 2000, 2018; Rosenbaum & Rubin, 1983; Wang & Blei, 2019). Put simply, this concept encapsulates the hope of the investigator that all relevant sources of variation have been adequately captured by the set of variables that are available for quantitative analysis. That is, assuming ignorability carries the bet by the analyst that all confounding covariates of no scientific interest are known and have been measured, and that there is no external knowledge outside of the data that would allow further characterizing the subject assessments: *y_diagnosis_* ⊥ *X_context of brain scan acquisition_* |*C*. In everyday research practice, this is an exceptionally strong assumption about the process that gave rise to the dataset at hand. This is because, if ignorability is truly satisfied, given the available covariates, the investigator is able to infer causal effects from the data without bias or skewing of the findings. In our present examination, however, propensity-score-matched subjects varied widely in the predictive performance and the extracted decision rules for classification. After repeated pulls of different population strata, the behavior of our predictive models diverged considerably when queried to distinguish subjects with a neuropsychiatric diagnosis and neurotypical controls. Even after deconfounding for subject age, sex at birth, and data acquisition site, our pairs of brain feature profile and target diagnosis were not mutually exchangeable and would also violate the independent and identically distributed (i.i.d.) assumption (exchangeability being a more lenient or more general assumption). This observation would be compatible with the more general conclusion that the ignorability assumption is not admissible in many or most brain-imaging studies given the types of confounding covariates that are measured in common large-scale datasets. It is plausible that there are lurking confounding variables, possibly attributable to latent population structure, that we are not yet in the habit of collecting or accounting for (Stürmer et al., 2010).

We admit several limitations about our quantitative findings and substantive conclusions. First, subjects were matched using propensity scores prior to conducting further steps of the analysis workflow. This decision resulted in smaller datasets for learning our predictive models, especially in the HBN cohort. To maximize the subjects available for model estimation, we used a caliper of 0.2 (commonly used in the literature (Austin, 2011b; Wang et al., 2013)), which, in turn, conceded to imperfect matches in certain cases. Here, smaller caliper values, such as for example 0.1, may have decreased the proportion of matches, but provided paired subjects with more similar covariates. Moreover, during stratification, subjects were grouped in equally-sized rather than strata that were equally-spaced in the diversity spectrum. We used equally-sized strata with the purpose of having the same number of subjects per strata. However, the covered diversity range may be different from one stratum to another, which may have impacted the computations. Finally, in our experiments, we found trends with different directionality when studying the impact of diversity on out-of-distribution performance in ABIDE and HBN. However, similar trends were found when only considering males or females in HBN. Therefore, our study showcases that sources of population stratification play an important role in predictions from brain imaging to a degree that is currently under-appreciated, independent of the directionality.

To conclude, we have added a sharp instrument to the analytical toolkit of neuroscience investigators. Diversity-aware machine learning pipelines allow tracking, unpacking and understanding *out-of-distribution generalization* in a variety of research agendas, such as studies involving brain scans. By bringing propensity scores to the neuroimaging community, subject stratification and cohort homogenization can be carried out in a seamless fashion with few or potentially many dozen covariates, be they categorical or continuous in nature. This principled solution can effectively detect and handle pivotal sources of demographic, clinical or technical variation, which can hurt the development of predictive models. Moreover, wider adoption of propensity scores can open new avenues to explore causal underpinnings of how brain representations are linked to the repertoire of human behavior. Based on our collective findings across two confederated neuroimaging efforts and different major psychiatric disorders, subject diversity is a key condition that determines generalization performance. Furthermore, the robustness of the predictive biomarkers cannot be guaranteed by the deconfounding practices that are the norm today.

## Acknowledgments

OB was funded by a Healthy Brains for Healthy Lives (HBHL) postdoctoral fellowship and is a member of the Quebec Autism Research Training (QART) program. BB acknowledges research support from the National Science and Engineering Research Council of Canada (NSERC Discovery-1304413), the Canadian Institutes of Health Research (CIHR FDN-154298), SickKids Foundation (NI17-039), Azrieli Center for Autism Research (ACAR-TACC), BrainCanada (Azrieli Future Leaders), and the Tier-2 Canada Research Chairs program. DB was supported by US NIH grant R01AG068563A and the Canadian Institutes of Health Research project grant 438531. DB was also supported by the Healthy Brains Healthy Lives initiative (Canada First Research Excellence fund), Google (Research Award, Teaching Award), and by the CIFAR Artificial Intelligence Chairs program (Canada Institute for Advanced Research).

## Author contributions

OB, DB and BB designed the experiments and wrote the manuscript. OB and DB led data analysis. CP, BP and VK aided with the experiments. SJH, RV, SZ, BTTY, ME, TG and JBP reviewed the manuscript. OB and DB are the corresponding authors of this work.

## Data availability statement

The imaging and phenotypic data were provided, in part, by the Autism Brain Imaging Data Exchange initiative (ABIDE-I and II; https://fcon_1000.projects.nitrc.org/indi/abide) and the Healthy Brain Network (https://fcon_1000.projects.nitrc.org/indi/cmi_healthy_brain_network). The specific subsets of data that were used in the present work are available from the authors upon request.

## Code availability statement

The code for running all experiments is available at https://github.com/OualidBenkarim/ps_diversity.

