## Supplementary Material for "The Cost of Untracked Diversity in Brain-Imaging Prediction"

### The Cost of Untracked Diversity in Brain-Imaging Prediction
### (Supplementary online material)

Oualid Benkarim[1], Casey Paquola[1,2], Bo-yong Park[1,3], Valeria Kebets[1], Seok-Jun Hong[4,5,6], Reinder Vos de Wael[1], Shaoshi Zhang[7,8,9], B.T. Thomas Yeo[7,8,9], Michael Eickenberg[10], Tian Ge[11], Jean-Baptiste Poline[1], Boris Bernhardt[1], Danilo Bzdok[1,12,13,14]

*[1]McConnell Brain Imaging Centre (BIC), Montreal Neurological Institute (MNI), Faculty of Medicine, McGill University, Montreal, Canada; [2]Institute of Neuroscience and Medicine (INM-1), Forschungszentrum Jülich, Jülich, Germany; [3]Department of Data Science, Inha University, South Korea; [4]Center for the Developing Brain, Child Mind Institute, New York, NY, USA; [5]Center for Neuroscience Imaging Research, Institute for Basic Science, Sungkyunkwan University, Suwon, South Korea; [6]Department of Biomedical Engineering, Sungkyunkwan University, Suwon, South Korea; [7]Department of Electrical and Computer Engineering, National University of Singapore, Singapore, Singapore; [8]Centre for Sleep and Cognition (CSC) & Centre for Translational Magnetic Resonance Research (TMR), National University of Singapore, Singapore, Singapore; [9]N.1 Institute for Health & Institute for Digital Medicine (WisDM), National University of Singapore, Singapore, Singapore; [10]Flatiron Institute, New York, NY, USA; [11]Psychiatric and Neurodevelopmental Genetics Unit, Center for Genomic Medicine, Massachusetts General Hospital, Boston, USA; [12]Department of Biomedical Engineering, Faculty of Medicine, McGill University, Montreal, Canada; [13]School of Computer Science, McGill University, Montreal, Canada; [14]Mila - Quebec Artificial Intelligence Institute, Montreal, Canada*

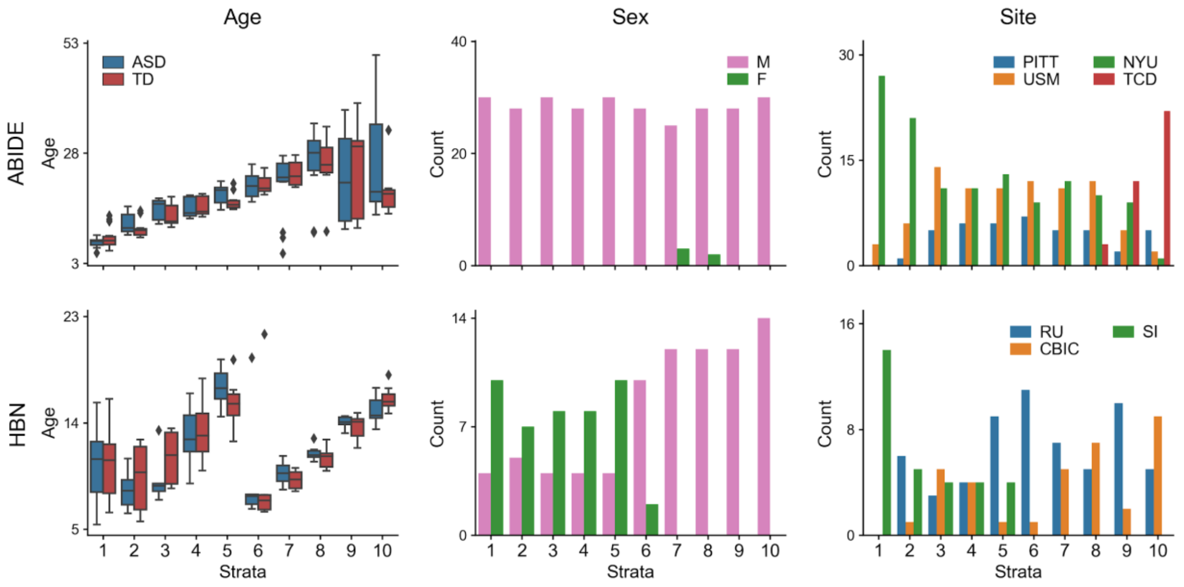


**Figure S1**. **Distribution of diversity indicators across strata**. From left to right, distribution of age, sex and site across strata in ABIDE (top) and HBN (bottom) datasets.

| | Site | Scanner | Modality | Sequence | TR (mm) | TE (mm) | TI (mm) | FA | Voxel (mm$^3$) |
|---|---|---|---|---|---|---|---|---|---|
| ABIDE | NYU | Siemens Allegra | T1w | 3D-TurboFLASH | 2530 | 3.25 | 1100 | 7° | 1.0×1.0×1.3 |
| | | | fMRI | 2D-EPI | 2000 | 15.00 | | 90° | 3.0×3.0×4.0 |
| | PITT | Siemens Allegra | T1w | 3D-MPRAGE | 2100 | 3.93 | 1000 | 7° | 1.1×1.1×1.1 |
| | | | fMRI | 2D-EPI | 1500 | 35.00 | | 70° | 3.1×3.1×4.0 |
| | TCD | Philips Achieva | T1w | 3D-MPRAGE | 3000 | 3.90 | 1150 | 8° | 0.9×0.9×0.9 |
| | | | fMRI | 2D-EPI | 2000 | 27.00 | | 90° | 3.0×3.0×3.2 |
| | USM | Siemens TrioTim | T1w | 3D-MPRAGE | 2300 | 2.91 | 900 | 9° | 1.0×1.0×1.2 |
| | | | fMRI | 2D-EPI | 2000 | 28.00 | | 90° | 3.4×3.4×3.0 |
| HBN | CIBIC | Siemens Prisma | fMRI | 2D-EPI | 800 | 30.00 | | 31° | 2.4×2.4×2.0 |
| | RU | Siemens TrioTim | fMRI | 2D-EPI | 800 | 30.00 | | 31° | 2.4×2.4×2.0 |
| | SI | Siemens Avanto | fMRI | 2D-EPI | 1450 | 40.00 | | 55° | 2.5×2.5×2.5 |

**Table S1**. **Scanner and data acquisition settings**. Abbreviations: TR, repetition time; TE, echo time; TI, inversion time; FA, flip angle; NYU, New York University Langone Medical Center; PITT, University of Pittsburgh, School of Medicine; TCD, Trinity Centre for Health Sciences, Trinity College Dublin; USM, University of Utah, School of Medicine; SI, Staten Island; RU, Rutgers University Brain Imaging Center; CBIC, CitiGroup Corcell Brain Imaging Center.

| | Site | N | | | | Sex (M/F) | Age |
|---|---|---|---|---|---|---|---|
| | | TD | ASD | ADHD | ANX | | |
| ABIDE | NYU | 70 | 56 | - | - | 121/5 | 15.0±7.4 |
| | PITT | 22 | 20 | - | - | 42/- | 20.2±7.1 |
| | TCD | 19 | 18 | - | - | 37/- | 15.2±3.3 |
| | USM | 40 | 52 | - | - | 92/- | 22.7±7.7 |
| HBN | CIBIC | 23 | 41 | 172 | 86 | 156/82 | 11.7±3.4 |
| | RU | 33 | 49 | 130 | 74 | 130/74 | 12.0±3.6 |
| | SI | 46 | 16 | 38 | 41 | 61/48 | 12.3±3.6 |

**Table S2**. **Demographics for each site.** Number of subjects (N), males/females (M/F) and mean age and standard deviation for each site. Note that some participants from HBN are diagnosed with more than one disorder. Abbreviations: NYU, New York University Langone Medical Center; PITT, University of Pittsburgh, School of Medicine; TCD, Trinity Centre for Health Sciences, Trinity College Dublin; USM, University of Utah, School of Medicine; SI, Staten Island; RU, Rutgers University Brain Imaging Center; CBIC, CitiGroup Corcell Brain Imaging Center.

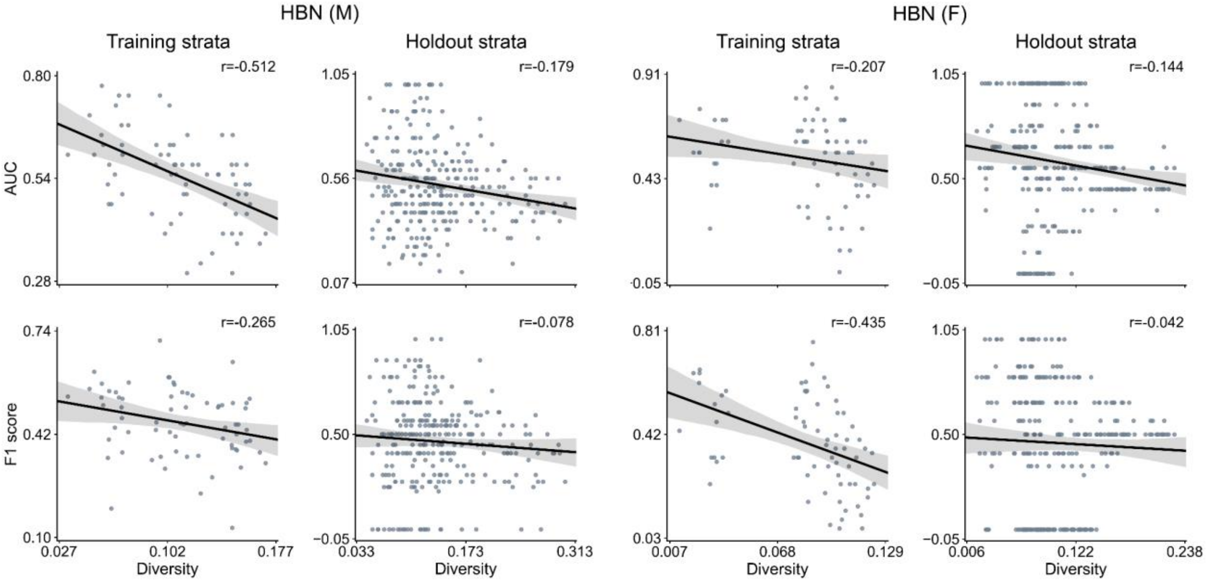


**Figure S2. Relationship between performance of predictive models and diversity in males and females (separately) in the HBN cohort.** Diversity is computed based on age and scanning site. After matching, analysis was carried out on 82 males and 44 females. Due to the small size of the data, we used all possible combinations of 4 out of 8 (instead of 5 out of 10) strata for training and the remaining 4 strata as holdout. Prediction accuracy based on area under the curve (AUC, top) and F1 score (bottom) in two subsets of the HBN dataset: only males (left) and only females (right). For each subset, the first column indicates the predictive model performance using a 10-fold cross-validation strategy based solely on the training set, where diversity is computed as the average of all pairwise absolute differences in propensity scores. The second column displays the performance for each single stratum in the holdout strata. Diversity denotes the mean absolute difference in propensity scores between the subjects of the training set and those in the held-out strata with unseen subjects. The strength of the association between performance and diversity is reported with Pearson's correlation coefficient (r).

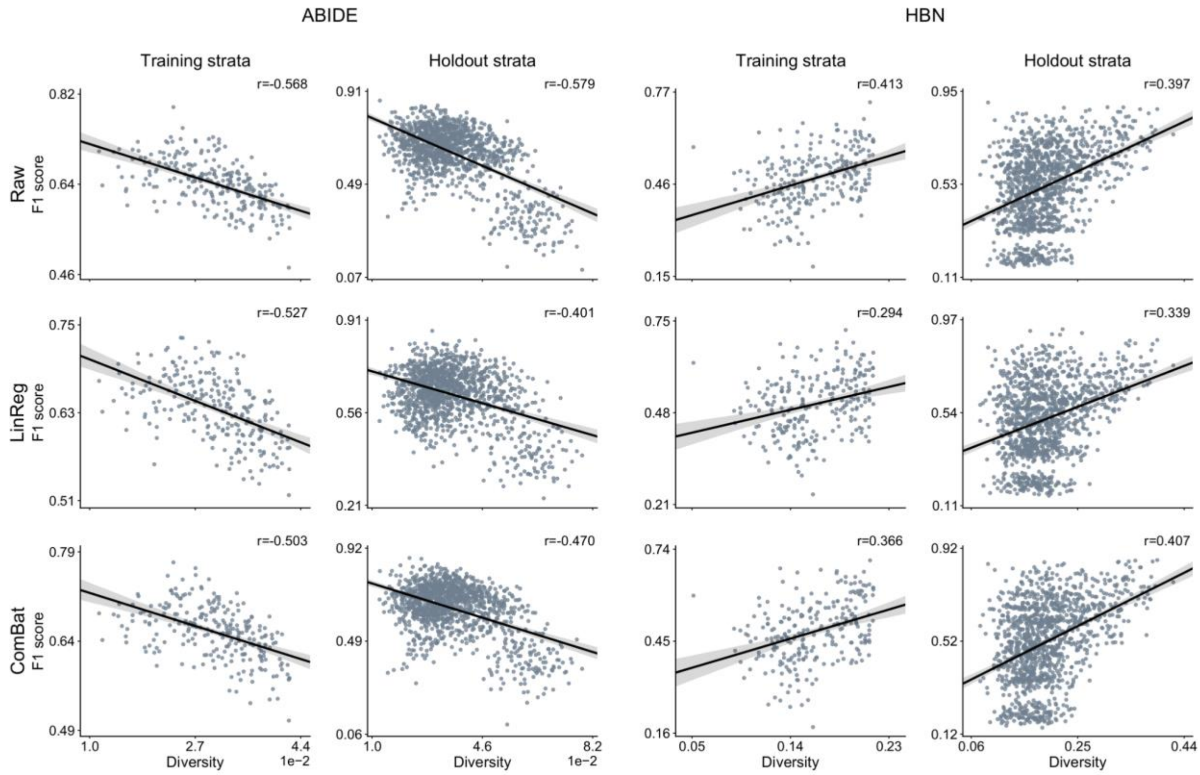


**Figure S3**. **Established and new deconfounding strategies appear insufficient to counteract escalating population diversity.** A means to quantify the behavior of predictive models is comparing its predictions on training and testing information. Performance is measured by F1 score on the original brain-imaging data (*Raw*, top), and after carrying out deconfounding steps on the brain feature prior to the pattern classification pipeline using standard linear-regression-based deconfounding (*LinReg*, middle) and recently proposed ComBat (bottom) approaches. Compared with the conventional nuisance removal using regression residuals, ComBat is a more advanced hierarchical regression to control for site differences. Results are reported for two different clinical cohorts: ABIDE (left) and HBN (right). For each cohort, the first column shows the model prediction performance using a 10-fold cross-validation (CV) strategy. Diversity is computed as the average of pairwise absolute differences in propensity scores between all subjects. Each dot is a cross-validated accuracy. The second column, for each cohort, displays the performance in the holdout subjects. Instead of reporting performance on all the subjects in the holdout data, here performance is assessed independently for subjects in each single stratum from the holdout data. Thus, diversity denotes the mean absolute difference in propensity scores between the subjects in the training set and those in the held-out stratum.

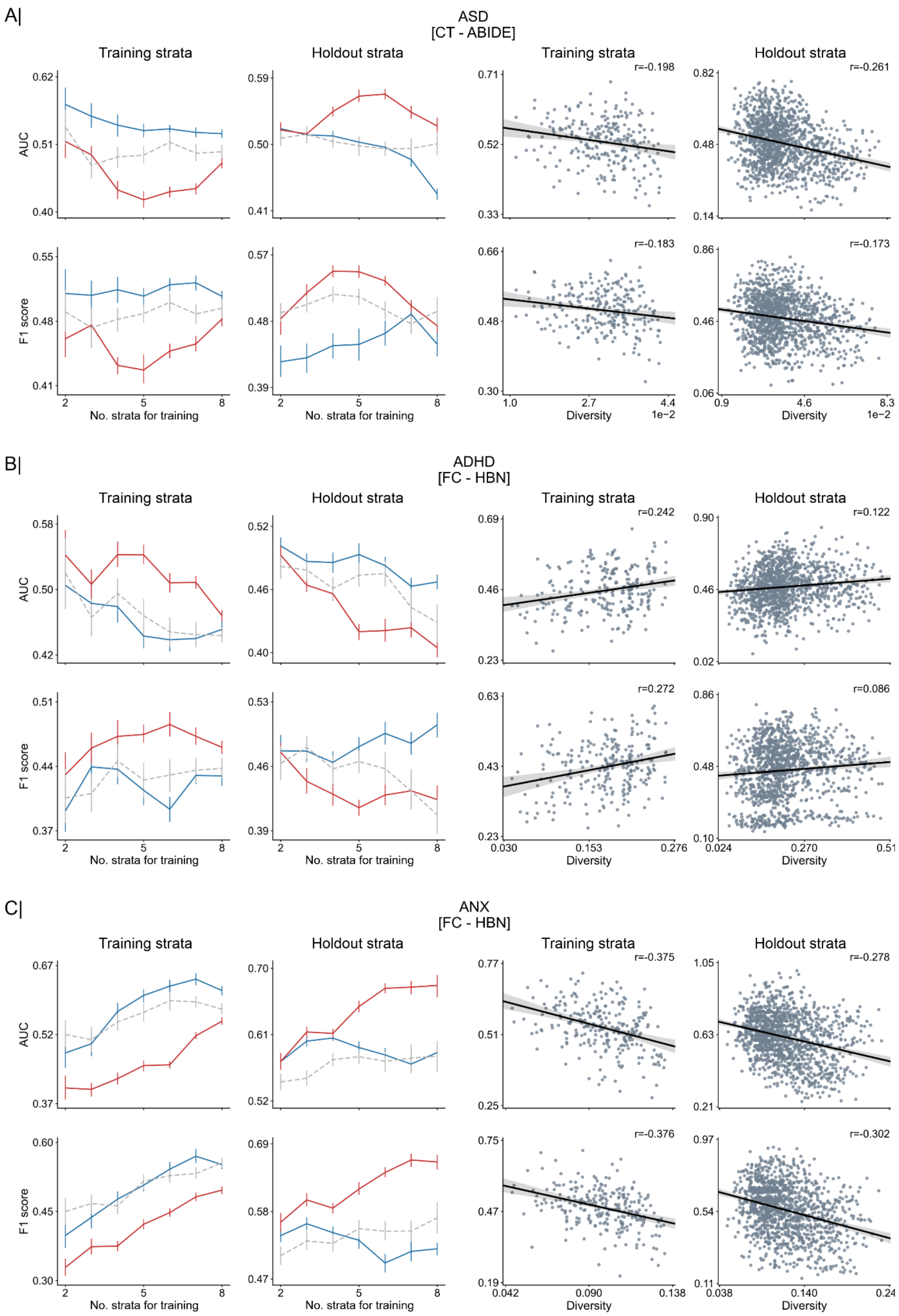


**Figure S4**. **Out-of-distribution prediction and performance of predictive models as a function of sample diversity.** Left: Comparison of model performance based on contiguous (Cont) and diverse (Div) training sets in the classification of ASD (top) using cortical thickness (CT) in ABIDE, and ADHD (middle) and ANX (bottom) based on functional connectivity in HBN. Performance is assessed using different sizes of training data (from 2 to 8 combined strata). For each disorder, the first column reports the performance using a 10-fold cross-validation strategy based solely on the training data, whereas the second column displays the performance in the holdout set, which contains the remaining subjects (from untouched strata). An additional model (i.e., Rand) is used as a baseline, where training subjects are randomly chosen regardless of their diversity (propensity scores). Right: Prediction performance based on all possible combinations of 5 out of 10 strata for training (and the remaining 5 for holdout). The first column reports the predictive model performance using a 10-fold cross-validation strategy based solely on the training set, where diversity is computed as the average of all pairwise absolute differences in propensity scores. The second column displays the performance for each single stratum in the holdout dataset, and diversity denotes the mean absolute difference in propensity scores between the subjects of the training set and those in the held-out subjects. The strength of the association between performance and diversity is reported with Pearson's correlation coefficient (r).

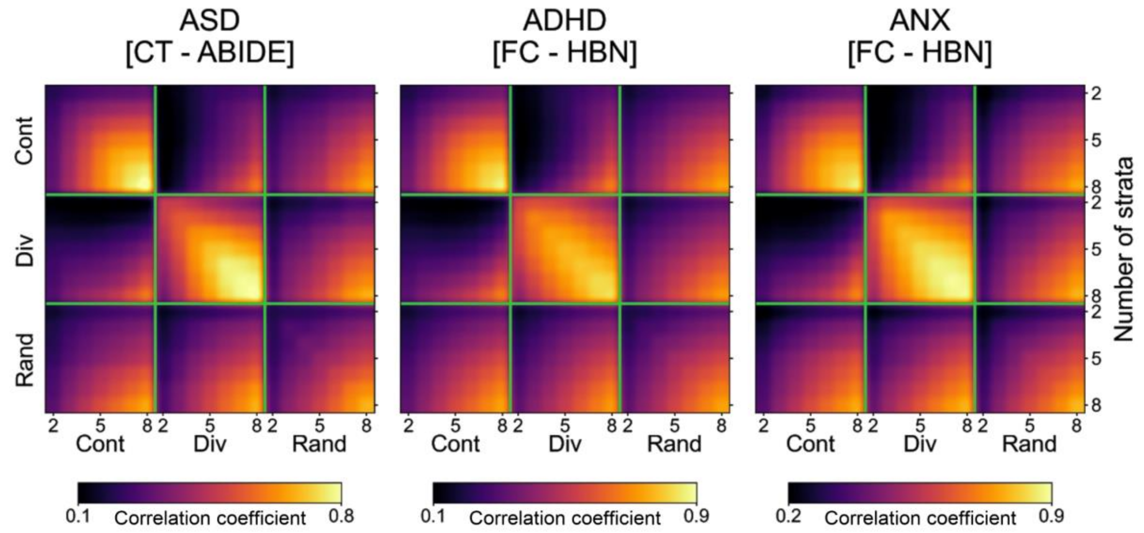


**Figure S5**. C**onsistency of extracted predictive patterns**. Consistency of model coefficients, as quantified by Pearson's correlation coefficient, obtained when using contiguous (Cont), diverse (Div) and random (Rand) sampling of subjects in the classification of autism using cortical thickness in the ABIDE dataset (left), and in the classification of ADHD (middle) and anxiety (right) based on functional connectivity in the HBN dataset. For each of these sampling schemes, consistency is shown for different numbers of combined strata used for predictive model training (from 2 to 8 combined strata), delineated by the green segments.

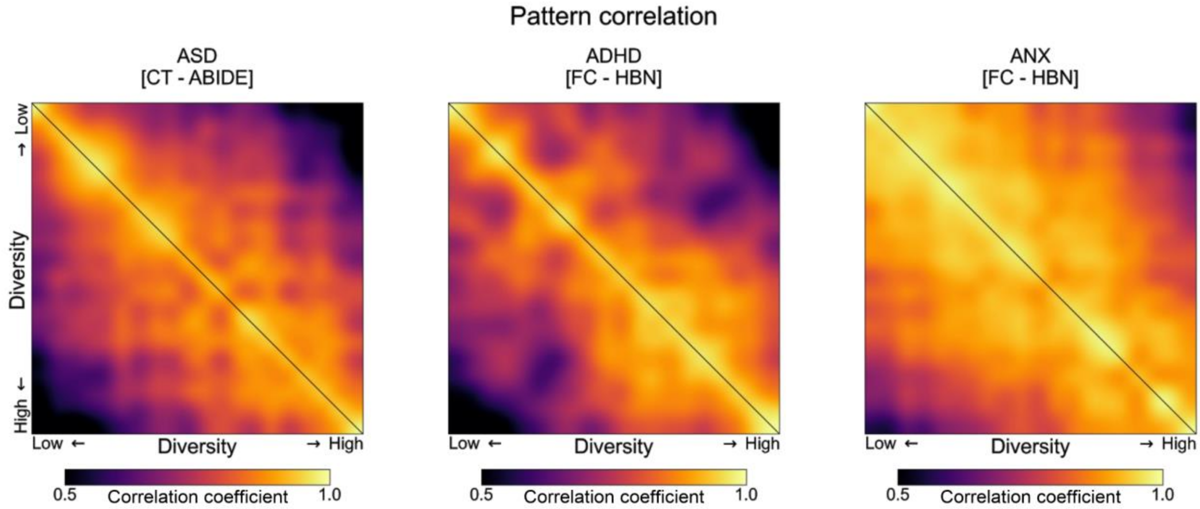


**Figure S6**. **Consistency of model-derived predictive patterns as a function of sample diversity**. Changes in predictive model coefficients with increasing diversity in the classification of autism using cortical thickness (CT) in the ABIDE dataset (left), and in the classification of ADHD (middle) and anxiety (right) based on functional connectivity in the HBN dataset. Consistency of model coefficients, in terms of Pearson's correlation, is obtained for each possible combination of training set (5 strata combined for training), where diversity is computed as the mean absolute difference in the propensity scores of the training observations. Pattern correlations are sorted according to their corresponding diversity (from low to high) based on the raw data (lower triangular part) and the ComBat-deconfounded data (upper part). Pearson correlation is based on the average of 10 consecutive model coefficients, according to their diversity.
